# A fungal pathogen effector that shapes host plant microbiota kills bacteria through lipoteichoic acid binding and membrane disruption

**DOI:** 10.64898/2026.05.26.727833

**Authors:** Gabriella C. Petti, Nick C. Snelders, Ashok K. Rout, Biwen Wang, Kathrin König, Tjalling Siersma, Jacob Biboy, Waldemar Vollmer, Leendert Hamoen, Jeroen R. Mesters, Daniel Friedrich, Alvaro Mallagaray, Bart P.H.J. Thomma

**Author notes:** These authors contributed equally to this work (Gabriella C. Petti, Nick C. Snelders).

## Abstract

Antimicrobial proteins are ancient and widespread molecules that contribute to survival across all domains of life. We previously showed that fungal plant pathogens secrete antimicrobial proteins to suppress antagonistic members of the host microbiota and thereby promote host colonization. The soil-borne plant pathogen *Verticillium dahliae* employs the bactericidal protein VdAve1 for this purpose. Here, we elucidate the mode of action of VdAve1 and define its mechanism as a distinct type of antimicrobial activity. Nuclear magnetic resonance (NMR) analysis revealed that VdAve1 adopts a jelly-roll barrel-like fold. Functionally, VdAve1 disrupts bacterial plasma membranes, and synthetic peptides derived from its positively charged regions retain antimicrobial activity, indicating that these regions contribute to its function. We further show that the bacterial model system *Bacillus subtilis* responds to VdAve1 by modifying teichoic acids, and that loss of these modifications increases bacterial sensitivity. Consistent with this, VdAve1 binds lipoteichoic acid (LTA), a major component of the Gram-positive cell wall. Together, our findings support a model in which VdAve1 binds LTA to localize at the bacterial surface, where it perturbs the plasma membrane, leading to membrane collapse and cell death.

**SIGNIFICANCE STATEMENT:** Plant-pathogenic microbes secrete effector proteins to promote host colonization. While these proteins are typically studied for their roles in host manipulation, some affect host-associated microbial communities by exhibiting antimicrobial activity. However, their modes of action remain largely unexplored. The fungal plant pathogen *Verticillium dahliae* secretes the antimicrobial effector VdAve1 to inhibit bacterial antagonists in the host microbiota. Here, we characterize the mode of action of VdAve1. We show that VdAve1 binds lipoteichoic acid, a major component of the cell wall of Gram-positive bacteria. This promotes VdAve1 accumulation close to the plasma membrane, and through electrostatic interactions it disrupts the plasma membrane. We further determine VdAve1 structure by NMR, revealing VdAve1 adopts a β-barrel-like fold. Together, our findings show the mechanism by which a fungal effector inhibits bacterial growth and thereby how it may influence host-associated bacterial communities.

## INTRODUCTION

Plants associate with complex and diverse microbiota that are highly dynamic, and whose composition is determined by various factors, including environmental conditions and plant genotype (1). Through the secretion of exudates, plants recruit beneficial microbes that enhance plant growth, nutrient acquisition, stress tolerance and disease resistance (1). Besides beneficial microbes, plant microbiota comprise commensal and pathogenic microbes. Under conducive circumstances, such as host immunity imbalances or abiotic stresses, pathogenic members of the plant microbiota and invading pathogenic microbes can cause disease that can be mitigated through competitive intermicrobial interactions, such as resource competition and secretion of antimicrobial compounds (2, 3). Additionally, microbes can stimulate plant immunity, enhance stress resilience and increase the capacity to withstand pathogen attack (1). Thus, the plant microbiota acts as an extended immune system (4).

Plant pathogens secrete effector proteins to facilitate host colonization. Initially thought to be solely involved in pathogen suppression of host immune responses, it is now recognized that effector proteins serve broader functions, such as nutrient acquisition (5). Given that plant-associated microbial communities contribute significantly to plant health, it is not surprising that growing evidence reveals that fungal pathogens secrete effector proteins with antimicrobial properties to selectively suppress microbial competitors during host, and more broadly, niche colonization (6). Seminal evidence comes from research on the soilborne fungus *Verticillium dahliae* that causes vascular wilt disease in a wide range of plant species, including crops (7). *V. dahliae secretes* several functionally characterized antimicrobial effector proteins during soil-dwelling, pathogenic and saprotrophic stages of its life-cycle to antagonize microbial niche competitors (8–11). Notably, the *V. dahliae* antimicrobial effector protein VdAve1 was shown to contribute to *V. dahliae* virulence by selectively suppressing bacterial antagonists in the host rhizosphere (8). Prior to this finding, however, VdAve1 was identified as an avirulence factor that is recognized by the tomato immune receptor Ve1, whereas it acts as a virulence factor that promotes disease development during infection on susceptible hosts lacking the Ve1 receptor (12). Interestingly, VdAve1 is homologous to a widespread class of plant peptides, called plant natriuretic peptides (PNPs) that are known to regulate water and ion homeostasis (13), and it has been suggested that VdAve1 was acquired by *V. dahliae* from plants via horizontal gene transfer (12). VdAve1 homologs are also found in few other phylogenetically unrelated fungal plant pathogens, such as *Colletotrichum higginsianum, Cercospora beticola* and *Fusarium oxysporum* (12), as well as in the bacterial plant pathogen *Xanthomonas citri* (14).

VdAve1 is a small (∼13 kDa) positively charged, cysteine-rich secreted protein that does not share sequence homology with known antimicrobial proteins (8). However, *in vitro* antimicrobial activity assays revealed selective antimicrobial activity. While all Gram-positive bacteria tested were inhibited, including the model Gram-positive bacterium *Bacillus subtilis*, Gram-negative bacteria showed differential sensitivity (8). Notably, scanning electron microscopy revealed that VdAve1 induces lysis of *B. subtilis* cells, indicating that VdAve1 has bactericidal activity (8). In this study we aimed to unravel the mode of action of VdAve1.

## RESULTS

### VdAve1 structure determination by NMR

To elucidate the mode of action of VdAve1, we first determined its structure by liquid-state NMR spectroscopy. Uniformly ^15^N- and ^15^N,^13^C-labelled mature VdAve1 was heterologously produced in *Escherichia coli* and purified. Because high protein concentrations can promote intermolecular interactions, concentration-dependent behaviour was assessed using ^1^H,^15^N-TRACT experiments (15)(16). The rotational correlation time (*τ*_*c*_) remained constant between 200–766 µM, indicating no detectable aggregation, although precipitation occurred at 766 µM upon prolonged incubation. Therefore, a concentration of 650 µM was used for subsequent NMR measurements.

Next, we acquired multidimensional NMR spectra for structure determination (Table S1), achieving 94% backbone amide assignment. Using the deep learning-based approach ARTINA (17), backbone assignment improved Cα coverage from 82% to 98%. ARTINA was then used to determine the structure yielding an ensemble of 20 conformers with a root mean square deviation of 2.22 Å (Figure 1b; Table S2). Validation via comparison of backbone torsion angles with TALOS-N predictions (18) showed good agreement (Table S3). Secondary structure analysis (model 1 of 20) indicated 7% alpha-helix, 23.2% beta-sheet, 13.4% turn and 56.4% coil. The structure features four anti-parallel β-strands (L2-T4, N110-Q115, Y49-C53 and M65-R73) forming a curved sheet, capped by a 7-residue alpha-helix (Q91-I97) and a small anti-parallel β-sheet (V32-V34, M87-L89). These six β-strands form two sheets resembling a jelly-roll fold/barrel, with alternative strand topology. Notably, the cysteines do not form disulphide bonds, consistent with unchanged HSQC spectra under reducing conditions (Figure S1). The overall surface charge is +6.21 (Figure 1c).

**Figure 1.**
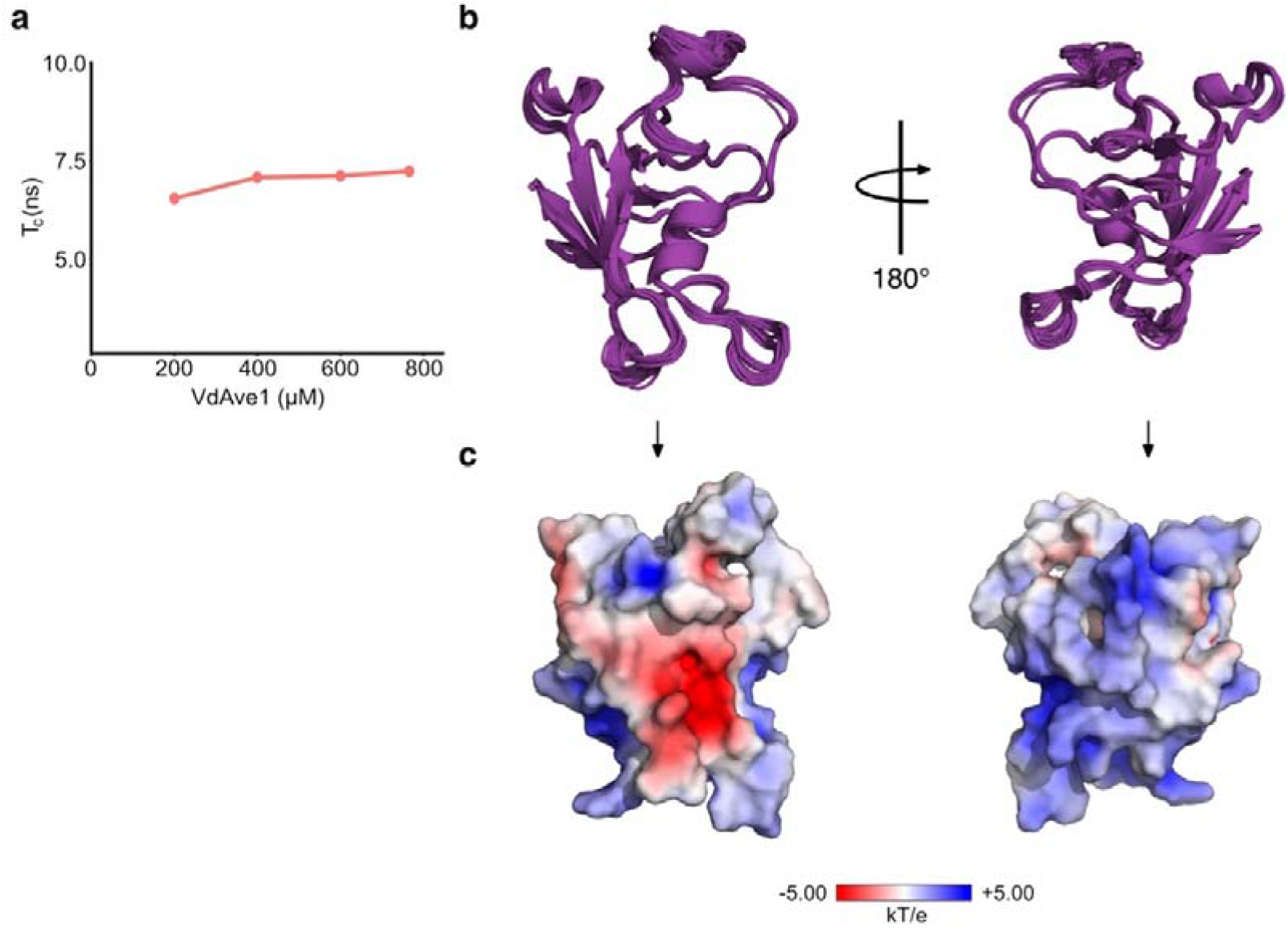
VdAve1 structure solved by liquid-state NMR spectroscopy. **a**. Rotational correlation times (*τ*_*c*_) calculated from ^1^H,^15^N TRACT experiments at different VdAve1 concentrations. **b**. Overlay of the 20 lowest-energy VdAve1 conformers determined by NMR spectroscopy. The structure ensemble has a root mean square deviation of 2.22 Å. The protein global fold comprises a six-stranded β-barrel flanked by an α-helix. **c**. VdAve1 surface charge distribution depicting positively charged (blue), negatively charged (red) and uncharged (white) regions of the protein.

To explore the mode of action of VdAve1, we performed a structural homology search, revealing homology to domain 1 (D1) of expansins, with top hits comprising expansins from *Clavibacter michiganensis, Gossypium hirsutum and Zea mays* (Table S4). Expansins are non-enzymatic plant proteins involved in cell wall loosening during development (19) and typically consist of two domains: D1 and D2, the latter mediating cell wall polysaccharide binding, while both are required for activity (20). The homology of VdAve1 to expansin D1 suggests a role in polysaccharide interactions.

### VdAve1 induces dissipation of *B. subtilis* cell membranes

Many cationic AMPs exert antimicrobial activity by targeting plasma membranes (21). Based on its net positive charge, we speculated that VdAve1 acts similarly. Membrane depolarization indicates membrane targeting, so we monitored localization of the cell division regulator protein MinD in *B. subtilis* upon incubation with VdAve1 (22). MinD associates with the plasma membrane at cell poles and division sites in a membrane potential-dependent manner (22, 23) and its localization via a MinD-GFP reporter is used as a proxy for depolarization (24). After 10 minutes of VdAve1 treatment, MinD-GFP delocalized from the cell poles and division sites (Figure 2a), whereas water or the unrelated *V. dahliae* effector protein Vd-D (25), had no effect (Figure 2a). These findings indicate that VdAve1 dissipates *B. subtilis* cell membranes.

**Figure 2.**
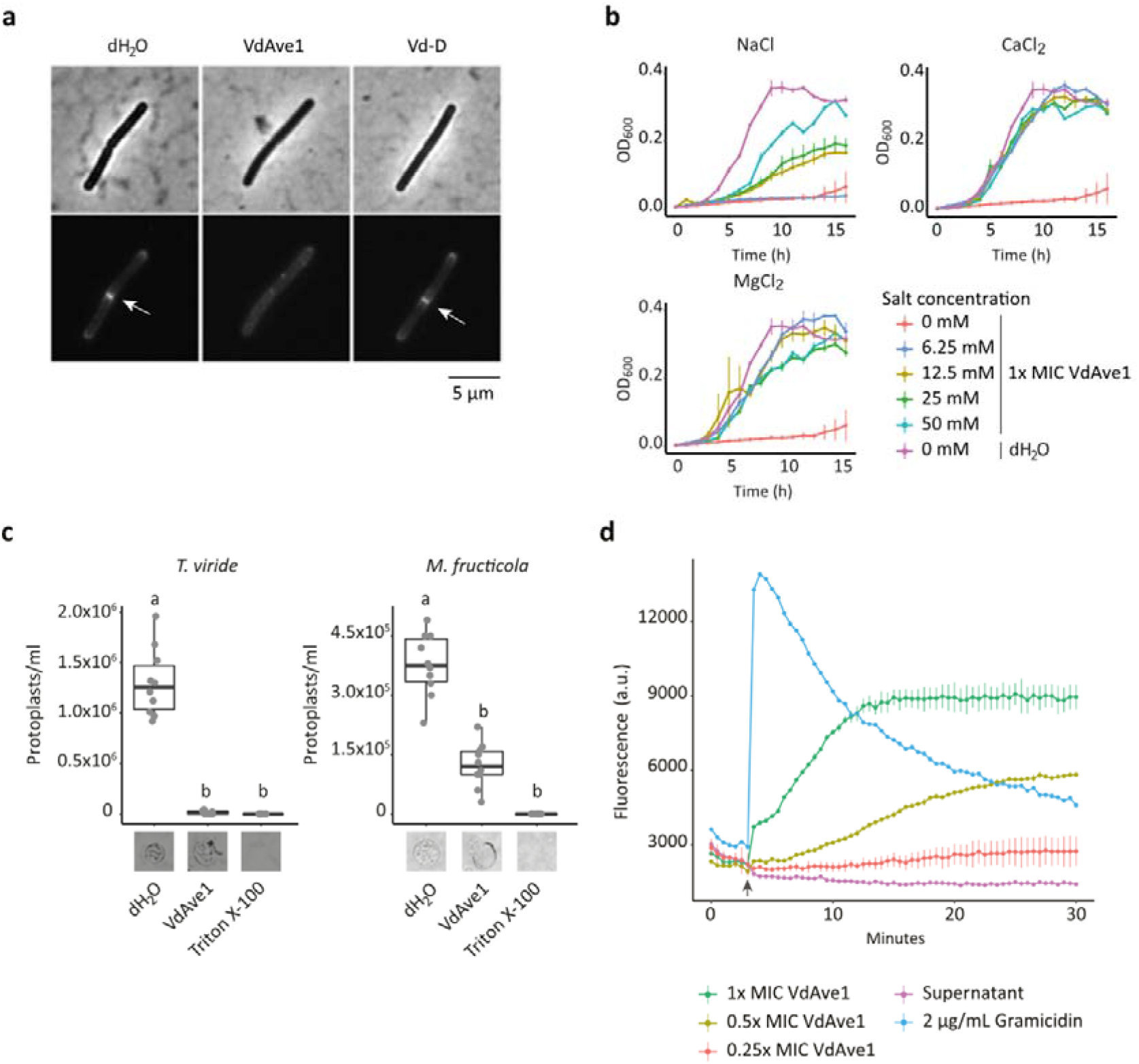
VdAve1 induces dissipation of *Bacillus subtilis* cell membranes. **a**. Cellular localization of the cell division regulator protein MinD-GFP in a *B. subtilis* reporter strain, upon 10 min of incubation with VdAve1, Vd-D or water. Treatment with 8 μM of VdAve1 induced delocalization of MinD-GFP from the cell poles and cell division sites. **b**. VdAve1 displays sensitivity to positively charged ions. VdAve1 activity on *B. subtilis* was tested in the presence of increasing NaCl, CaCl_2_ and MgCl_2_ concentrations. Growth curves represent mean OD_600_ values ± standard deviation (n=3). **c**. VdAve1 causes lysis of protoplasts from the fungal species *Trichoderma viride* and *Monilinia fructicola*. Triton X-100 was used as positive control. Representative pictures of the corresponding protoplasts are displayed under the boxplots. Different letter labels represent statistically significant differences according to one-way ANOVA and Tukey’s post-hoc test; p<0.0001, N=10. **d**. VdAve1 induces gradual depolarization of *B. subtilis* cell membranes. Release of the fluorescence potentiometric probe DiSC_3_(5) was monitored over time as means of membrane depolarization after incubation with different VdAve1 concentrations or the pore-forming peptides gramicidin. The arrow indicates the time point of protein administration. Graphs represent mean values ± standard deviation (n=3).

Cationic AMPs often rely on electrostatic interactions with negatively charged cell envelope components and therefore are often salt sensitive (21). Given its positive charge, we hypothesized that VdAve1 activity similarly requires electrostatic interactions. To test this, we monitored *B. subtilis* growth inhibition by VdAve1 in the presence of various concentrations of salts. Interestingly, VdAve1 activity was reduced by Na^+^ and completely abolished by ≥6.25 mM Mg^2+^ or Ca^2+^ (Figure 2b), indicating strong cation sensitivity and suggesting electrostatic interactions with bacterial surface components.

To test whether VdAve1 directly targets membranes, we examined fungal protoplasts. Although bacterial and fungal membranes differ, many membrane-active AMPs affect both (26). Notably, VdAve1 caused lysis of *Trichoderma viride* and *Monilinia fructicola* protoplasts (Figure 2c), indicating that VdAve1 displays membrane-lytic activity.

Many cationic AMPs disrupt membranes through the formation of pores, resulting in rapid depolarization (21). To further study the effect of VdAve1 on plasma membranes, we measured membrane depolarization kinetics in *B. subtilis* using the potentiometric dye DiSC_3_(5) (27). As expected, the pore-forming control peptides gramicidin ABC induced immediate depolarization (Figure 2d) (28). In contrast, VdAve1 induced a gradual membrane depolarization (Figure 2d), suggesting that membrane disruption is not achieved through pore formation.

### VdAve1-derived cationic peptides display antimicrobial activity

To investigate whether specific regions of VdAve1 are responsible for *B. subtilis* membrane disruption, we designed partially overlapping 20 amino acid-long VdAve1-derived peptides, collectively spanning the mature protein sequence (Table S5), and screened them for antimicrobial activity. Strikingly, two peptides, aa41-60 and aa101-116, completely inhibited *B. subtilis* growth (Figure 3a). Interestingly, these peptides have the highest predicted positive charge (+4.91 for aa41-60, +3.09 for aa101-116) and correspond to adjacent, surface-exposed antiparallel β-strands in the VdAve1 structure (Figure 3b), suggesting that this region directly contributes to membrane perturbation.

**Figure 3.**
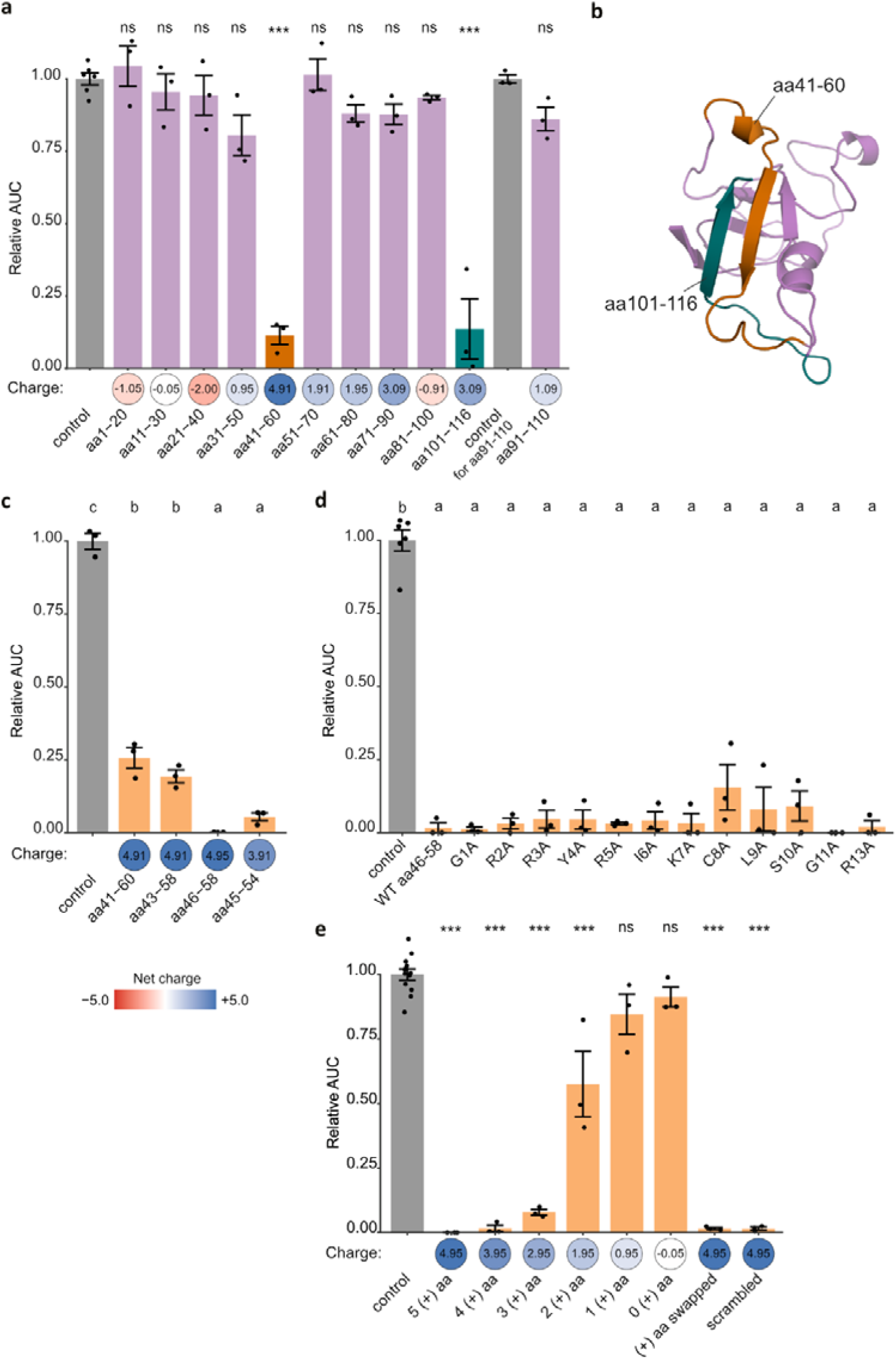
VdAve1-derived cationic peptides display antimicrobial activity. **a**. Antimicrobial activity test of 20 aa long partially overlapping peptides that collectively span the mature VdAve1 sequence against *B. subtilis*. Peptides aa41-60 (orange bar) and aa101-116 (petrol bar) significantly reduced *B. subtilis* growth. The plot shows area under the growth curve (AUC) of *B. subtilis* treated with the respective peptide relative to the corresponding control. Bars indicate means, error bars represent standard errors, individual data points are shown (n=6 for the control, n=3 for all others). Statistical significance was assessed by one-way ANOVA and post-hoc Tukey HSD test (\*\*\**P*⍰≤0.001; ns, not significant). Charges of corresponding peptides are displayed below the bar plot. **b**. VdAve1 structure indicating the regions corresponding to the antimicrobial peptides: aa41-60 (orange) and aa101-116 (petrol). **c**. Antimicrobial activity test of truncated variants of peptide aa41-60. Plot shows area under the growth curve (AUC) of *B. subtilis* treated with the respective peptide relative to the control. Bars indicate means, error bars represent standard errors, individual data points are shown (n=3). Letters indicate significant differences based on one-way ANOVA and post-hoc Tukey HSD test. Charges of corresponding peptides are displayed below the bar plot. **d**. Antimicrobial activity test of single amino acid mutants of peptide aa46-58. The plot shows area under the growth curve (AUC) of *B. subtilis* treated with the respective peptide relative to the control treatment. Bars indicate mean, error bars represent standard error, individual data points are shown (n=6 for the control, n=3 for all others). Different letter labels indicate significant differences based on one-way ANOVA and post-hoc Tukey HSD test. **e**. Antimicrobial activity test of aa46-58 mutated in increasing number of positively charged residues as well as a randomized version of the peptide. The plot shows area under the growth curve (AUC) of *B. subtilis* treated with the respective peptide relative to the control treatment. Bars indicate mean, error bars represent standard error, individual data points are shown (n=12 for the control, n=3 for all others). Statistical significance was assessed by one-way ANOVA and post-hoc Tukey HSD test (\*\*\**P*⍰≤0.001; \*\**P*⍰≤0.001; \**P*⍰≤0.05; ns, not significant). Charges of corresponding peptides are displayed below the bar plot.

Next, we further characterized the antimicrobial activity of peptide aa41-60, showing the highest charge. Truncation analysis showed that the peptide antimicrobial activity correlated with charge and length, with aa46-58 (13 aa) showing the strongest inhibition (Table S5, Figure 3c). Single residue mutations within this peptide did not reduce its activity (Table S5, Figure 3d), indicating that the antimicrobial activity cannot be assigned to a particular amino acid.

To test the role of charge in aa46-58 activity, positively charged residues were progressively replaced with alanines, resulting in peptides with positively charged residues ranging from four to zero. Antimicrobial activity decreased with decreasing charge and was lost when four of five charged residues were mutated (Figure 3e). Interestingly, swapping arginines with lysines, and *vice versa*, did not affect peptide antimicrobial activity (Table S5; Fig. 3d), indicating that the activity depends on the presence of positively charged residues, irrespective of their side chains.

To test whether peptide aa46-58 activity is sequence-dependent, we randomized its amino acid order while keeping length and composition unchanged (Table S5). Interestingly, the scrambled peptide completely inhibited bacterial growth (Figure 3d), suggesting that the activity does not depend on the exact amino acid sequence, but rather on overall physicochemical properties. Additionally, it suggests that the antimicrobial activity of the peptide does not rely on sequence-specific interactions, like receptor-binding. Overall, our findings suggest that the antimicrobial activity of aa46-58 depends on its net charge.

### Antimicrobial activity of the VdAve1-derived peptide against a Gram-negative bacterium

It was previously shown that VdAve1 displays antimicrobial activity against particular Gram-negative bacteria (8). To investigate whether the activity of the peptide extends to Gram-negative bacteria, we tested aa46-58 and its single amino acid mutants against *Sphingopyxis macrogoltabida* (Figure S2; Table S5). In contrast to what was observed with *B. subtilis*, substitution of the cysteine with alanine (C8A) completely impaired the peptide activity. Replacement with serine (C8S), which preserves polarity but lacks a thiol group, similarly resulted in loss of activity, indicating that the cysteine thiol is required for the antimicrobial activity. To assess the role of the peptide charge for antimicrobial activity, we tested the ability of the aa46-58 mutant lacking positively charged amino acids to inhibit *S. macrogoltabida* growth. Interestingly, the antimicrobial activity was completely lost (Figure S2), indicating that the peptide activity depends on its charge also against Gram-negative bacteria. Furthermore, the scrambled mutant of aa46-58 retained full activity (Figure S2, Table S5), indicating that the activity of the peptide does not dependent on its amino acid sequence and that the contribution of the cysteine is position-independent. Together, our findings indicate that the antimicrobial activity of aa46-58 against *S. macrogoltabida* depends on its charge and the presence of a cysteine, possibly required for the formation of disulphide-linked peptide dimers with increased molecular charge.

### Identification of genes involved in the response and resistance to VdAve1

To improve our understanding of the mode of action of VdAve1, we examined transcriptional changes of *B. subtilis* exposed to sub-minimum inhibitory VdAve1 concentrations (0.5x MIC) (8), after 5 and 20 minutes of incubation. The most strongly induced genes in response to VdAve1 at both time points were the *sigV* operon genes: *sigV*, which encodes for the sigma factor *σ*^V^, *rsiV, oatA* and *yrhK* (Table S6). *σ*^V^ regulates genes involved in peptidoglycan modifications and it was previously implicated in resistance to lysozyme (29). Besides its own operon, *σ*^V^ induces the *dlt* operon, which mediates D-alanylation of teichoic acids (29). Accordingly, all *dlt* operon genes were induced in response to VdAve1 (Table S7). Moreover, exposure to VdAve1 led to the induction of genes involved in cation export, such as *ydbO, cadA, ykkD* and *nhaK* (30–33) (Table S7). Cation export might hinder electrostatic interactions between VdAve1 and its potential negatively charged target, in a similar fashion as observed when testing VdAve1 activity in the presence of salts (Figure 2b). Finally, the phospholipid desaturase *des*, associated with increased membrane fluidity (34), was downregulated in response to VdAve1, suggesting that VdAve1 impacts membrane fluidity.

To further substantiate the involvement of the differentially expressed genes in protection to VdAve1, and find a possible molecular binding target, we screened and sequenced a *B. subtilis* transposon insertion library under VdAve1 exposure over nine generations. Consistent with the transcriptome analysis, disruption of *σ*^V^-regulated genes, *oatA* and *dltA–E*, increased sensitivity to VdAve1, while loss of *rsiV*, which regulates *σ*^V^ activity (35), improved survivability in the presence of VdAve1 (Table S8). Mutations in genes associated with teichoic acid synthesis or modification (*galE, gtaB, ugtP*) (36–38) and in cell wall-related enzymes (*pbpX, cwlO, dacA*) (39–41) also increased sensitivity (Table S8). Conversely, disruption of *rsiX*, the regulator of *σ*^X^ (42), increased tolerance to VdAve1. Interestingly, *σ*^X^ was previously shown to contribute to resistance to cationic antimicrobial peptides by regulating cell surface modifications through the activation of other operons, including the *dlt* operon (43). Even though we could not identify a specific molecular target at the cell envelope, the transposon screen corroborated the relevance of genes involved in cell envelope modification for tolerance of *B. subtilis* to VdAve1 treatment.

### VdAve1 does not hydrolyse peptidoglycan

Transcriptome profiling and transposon sequencing identified the *SigV* operon, previously implicated in lysozyme resistance (29), as a key component of *B. subtilis* response to VdAve1. Additionally, the bacterial response to VdAve1 involved genes associated with teichoic acid modification and biosynthesis, which are also known to contribute to lysozyme resistance (29). These parallels suggested that VdAve1 may function as a lysozyme. To test this, we incubated peptidoglycan from *B. subtilis* with VdAve1 in two buffer conditions, including a low-salt one to account for VdAve1 salt sensitivity, and performed high-performance liquid chromatography (HPLC) to detect possible products released. In both conditions, the elution profile did not show distinct peaks corresponding to muropeptides, indicating that VdAve1 does not digest peptidoglycan (Figure S3). In contrast, digestion with cellosyl, a muramidase from *Streptomyces coelicolor* (44), produced the expected muropeptide profile, which was unchanged upon co-incubation with VdAve1 (Figure S3). These results indicate that VdAve1 does not exhibit lysozyme(-like) activity on peptidoglycan and does not digest muropeptides.

### VdAve1 binds lipoteichoic acid

The transcriptomic and transposon screening analysis highlighted the importance of peptidoglycan and teichoic acid modifications for *B. subtilis* tolerance to VdAve1. To evaluate possible interactions of VdAve1 with these molecules, we performed ^1^H,^15^N HSQC-based chemical shift perturbation (CSP) experiments. No CSPs were observed upon exposure to *B. subtilis* peptidoglycan, indicating no binding, whereas *B. subtilis* lipoteichoic acid induced clear CSPs (Figure S4), suggesting that VdAve1 binds LTA. LTA titration (0.2-0.6 mg/mL) revealed concentration-dependent CSPs in specific residues, such as K84 and A85 (Figure 4a, expanded view). Next, we quantified CSPs (Δ*δ*_NH_) by comparing backbone amide shifts in the absence and presence of 0.6 mg/mL LTA (Figure 4b) and mapped these onto the VdAve1 structure (Figure 4c). The perturbations localized to one side of the protein, forming two regions characterized by large (Δ*δ*_NH_>2*σ*) and minor CSPs (Δ*δ*_NH_>*σ*,). K84 displayed the largest perturbation, suggesting a potential involvement in LTA binding. Minor CSPs were also detected in the His-tag used for affinity purification (Figure S5), likely caused by unspecific interactions or allosteric effects due to LTA binding.

**Figure 4.**
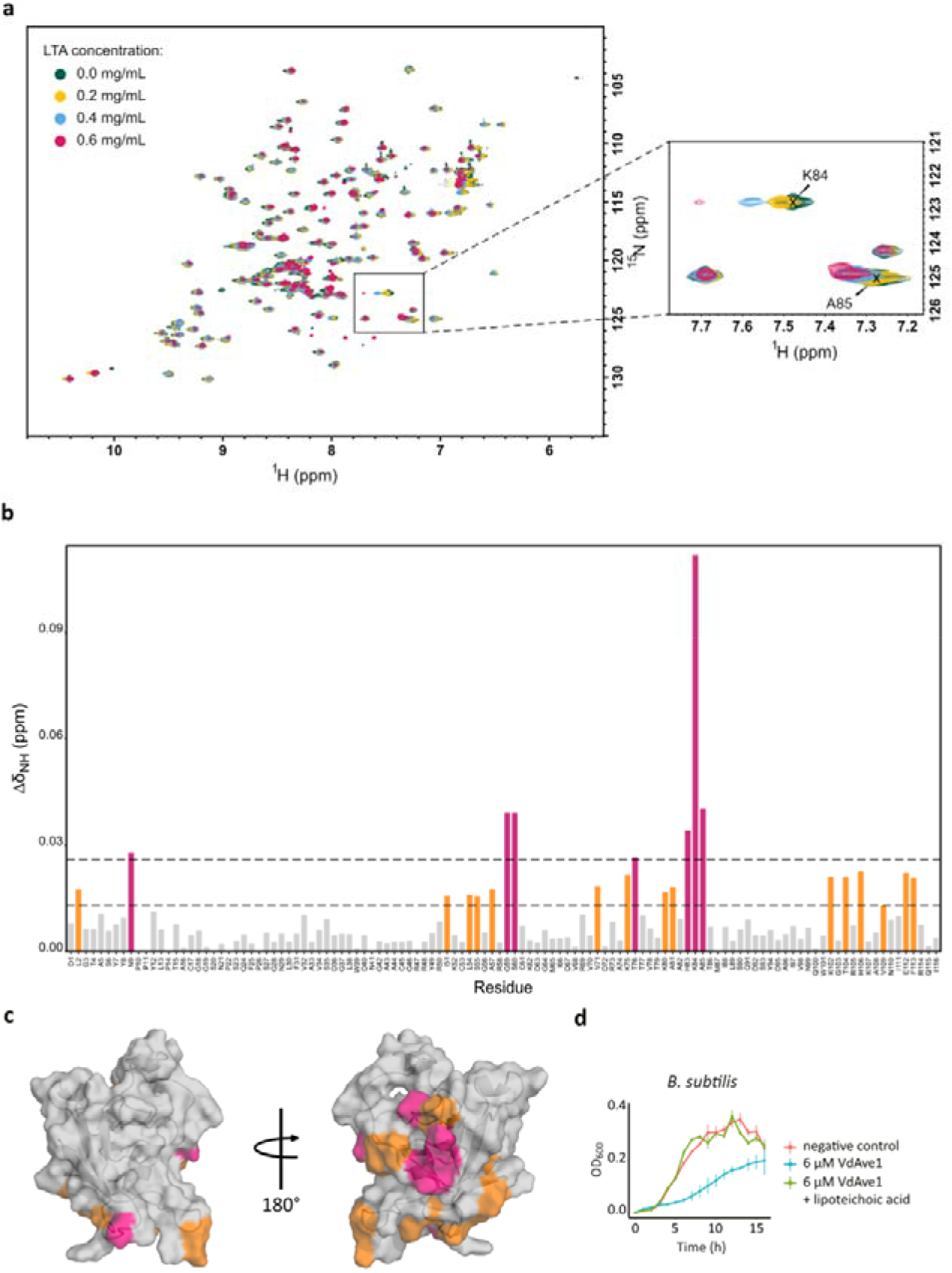
VdAve1 binds lipoteichoic acid. **a**.^1^H, ^15^N HSQC spectra of VdAve1 upon binding with increasing concentrations of lipoteichoic acid (LTA): 0.0 mg/mL (green), 0.2 mg/mL (yellow), 0.4 mg/mL (cyan), 0.6 mg/mL (pink). Full spectra (left) and the expanded view (right) show peaks perturbations. **b**. Chemical shift perturbations (CSPs) for each VdAve1 residue induced by 0.6 mg/mL LTA. The Δ*δ*_NH_ values were color-coded, Δ*δ*_NH_>2*σ* pink and Δ*δ*_NH_>*σ* orange. **c**. CSPs mapped onto the VdAve1 structure. **d**. Preincubation VdAve1 with an equal concentration of LTA from *B. subtilis* inhibits VdAve1 activity. Curves represent mean values ± standard deviation (n=3).

To investigate the relevance of LTA binding for the mode of action of VdAve1, we preincubated VdAve1 with *B. subtilis* LTA prior to antimicrobial assays with *B. subtilis* (Figure 4d). Interestingly, preincubation abolished VdAve1 activity, indicating that binding to LTA prevents interaction of the protein with the bacterial cell envelope, thereby impairing its ability to inhibit bacterial growth.

Because Gram-negative bacteria lack LTA, but possess the glycolipid lipopolysaccharide (LPS), we speculated that LPS could serve as an alternative binding target for VdAve1. However, LPS from *Pseudomonas aeruginosa* did not induce CSPs (Figure S6), suggesting that VdAve1 does not bind LPS.

## DISCUSSION

Antimicrobial proteins (AMPs) are ancient, widely distributed defence molecules across all domains of life (21). Despite their diversity, AMPs are generally small, cationic molecules with broad-spectrum activity. Their antimicrobial activity commonly arises from electrostatic interactions with negatively charged bacterial membranes, leading to membrane destabilization and cell death (45). Membrane disruption is generally described by two models: pore formation via membrane insertion, or surface accumulation that disrupts membrane integrity (21).

We previously showed that the soil-borne plant pathogen *V. dahliae* exploits the bactericidal effector protein VdAve1 to suppress bacterial antagonists during host colonization (8). Here, we show that VdAve1 induces dissipation of the *B. subtilis* plasma membrane, leading to cell leakage and bacterial death. This activity depends on electrostatic interactions, as evidenced by the salt sensitivity of the antimicrobial activity and the requirement for positively charged residues in active VdAve1-derived peptides.

Using independent approaches, we found that genes involved in cell wall modification are induced in response to VdAve1 and affect *B. subtilis* tolerance to VdAve1. Particularly, D-alanylation of teichoic acid was an important contributor to this tolerance.

Teichoic acids are anionic molecules in Gram-positive cell walls, present either anchored to the plasma membrane as lipoteichoic acid (LTA) or connected to the cell wall as wall teichoic acid (46). D-alanylation reduces their negative charge, decreasing the electrostatic attraction of cationic AMPs to the bacterial surface (47). The involvement of teichoic acid modification in both the response and tolerance to VdAve1 suggested that these molecules may serve as direct targets. Consistent with this, we show that VdAve1 binds purified LTA.

Membrane-disrupting AMPs are proposed to interact with cell wall components and precursors prior to membrane disruption (48). For instance, nisin binds lipid II, a peptidoglycan precursor, prior to membrane disruption (49). Lipid II binding, besides inhibiting cell wall biosynthesis, aids nisin-mediated membrane permeabilization (49). Lipid II is anchored in the plasma membrane, thereby localizing AMPs at the membrane interface and promoting their insertion into the bacterial membrane. Similarly, LTA has been proposed as a potential AMP interaction site (48). Several membrane-disrupting AMPs have been suggested to interact with LTA, as exogenous LTA reduced AMP activity, although direct binding evidence remains limited (50, 51).

Importantly, our data suggest that LTA binding and membrane disruption involve distinct regions of VdAve1. The NMR analyses did not reveal major conformational changes upon LTA binding, arguing against a mechanism in which LTA interaction activates or exposes a membrane-targeting domain. Instead, we propose that LTA binding functions primarily to localize or retain VdAve1 at the bacterial surface, thereby facilitating interaction of the distinct membrane-active region with the cell membrane, ultimately leading to membrane dissipation and cell death.

We previously showed that *V. dahliae* uses VdAve1 to suppress the growth of Gram-negative Sphingomonads during host colonization (8). Although Gram-negative bacteria lack LTA, they possess lipopolysaccharide (LPS), which shares structural and functional similarities with LTA (46, 52), thus representing a plausible alternative binding target for VdAve1. However, VdAve1 does not interact with LPS from *P. aeruginosa*, indicating that its activity against Gram-negative bacteria does not require a cell wall target or involves a different one. Notably, activity of VdAve1-derived peptides against Gram-negative bacteria requires a cysteine residue in addition to charge-based interactions. This residue may be necessary for the formation of peptide dimers, enhancing their overall charge and structural stability.

However, it remains unclear why this feature is not required for targeting Gram-positive bacteria.

By solving the VdAve1 structure, we present an experimentally determined structure of a PNP-like protein, providing a foundation for understanding the biological function of this underexplored protein family. The VdAve1 fold closely resembles domain 1 of expansins, proteins known to mediate plant cell wall loosening (19). Although expansins are not known to possess antimicrobial activity, the shared fold suggests that related structural features support interactions with microbial cell surfaces, potentially influencing bacterial envelope integrity.

Structural analysis revealed that VdAve1 lacks disulphide bridges, which are commonly found in effector proteins and are thought to confer stability in protease-rich environments such as the host apoplast (53). Despite lacking intramolecular disulphide bonds, VdAve1 is functionally active, as evidenced by its antimicrobial activity, indicating that the reduced state of its cysteine residues does not compromise its biological function. Instead, the presence of reduced cysteines raises the possibility that they might play a role in VdAve1 activity. Notably, cysteine is required for the activity of VdAve1-derived peptides against Gram-negative bacteria, suggesting that reduced cysteines in the full-length protein enable the formation of intermolecular disulphide bonds between VdAve1 molecules at the bacterial surface. Alternatively, proteolytic processing by host proteases may generate cysteine-containing peptides that become active against Gram-negative bacteria upon disulphide-mediated oligomerization. Another possibility is that the free thiol groups participate in metal ion coordination, which has been shown to enhance the antimicrobial activity of several AMPs (54).

In conclusion, our findings demonstrate that VdAve1 inhibits microbial growth by disrupting plasma membranes through electrostatic interactions. Additionally, we show that VdAve1 binds LTA in the cell wall of Gram-positive bacteria, likely serving as a docking platform that facilitates its accumulation at the bacterial surface before membrane destabilization. These findings establish how a fungal effector protein directly suppresses bacterial competitors during host colonization.

## MATERIALS AND METHODS

### Expression and purification of recombinant VdAve1

Recombinant VdAve1 was expressed in *Escherichia coli* BL21 cells carrying pET15b-VdAve1 and purified from inclusion bodies followed by refolding as previously described (8). Proteins were stored in 30 mM potassium phosphate buffer (pH 6.5). Detailed procedures are provided in SI Materials and Methods.

### NMR spectroscopy and structure determination

NMR experiments were performed at 298 K on Bruker Avance III HD spectrometers operating at 600 or 500 MHz equipped with cryogenic probes. Backbone and side-chain resonances were assigned using standard triple-resonance and side-chain experiments. Distance restraints for structure calculation were derived from ^1^H-^15^N and ^1^H-^13^C NOESY-HSQC spectra, and structures were determined using automated assignment and structure calculation with ARTINA (17). Samples for structure determination contained 600-650 µM ^13^C/^15^N-labeled VdAve1 in phosphate buffer (pH 6.6), and protein concentrations ranging from ∼150 to 750 µM were used for relaxation and titration experiments (see SI Materials and Methods for details).

### Chemical shift perturbation (CSP) NMR experiments

^1^H-^15^N HSQC spectra for CSP NMR experiments were collected in the presence of 1 mg/mL of *B. subtilis* peptidoglycan, and 0.2, 0.4, 0.6, 1.0 mg/mL of *B. subtilis* lipoteichoic acid (LTA) (Sigma-Aldrich, St. Louis, USA). CSPs were calculated based on 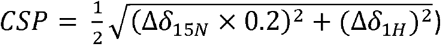 and are listed in Table S9. CSPs greater than 2*σ* were considered significant and further interpreted as signals that reflect interacting residues.

### MinD delocalization assay

MinD delocalization was conducted as previously described (24) with minor modifications. *B. subtilis* strain TB35, expressing MinD-GFP, was cultivated in 0.5× YT medium supplemented with 0.1% xylose at 30°C until reaching an OD_600_ of 0.1. Cultures were treated with 8 μM (1× MIC) VdAve1, Vd-D or sterile ultrapure water for 10 minutes, immobilized on 1% agarose thin films. MinD-GFP localization was analysed using an Olympus BX 50 microscope equipped with a Photometrics CoolSNAP fx digital camera.

### Activity assays of VdAve1 and VdAve1-derived peptides on bacteria

B. subtilis strain 168 was grown on lysogeny broth agar (LBA) at 28⍰°C. A single colony was used to inoculate overnight cultures in low salt LB (10 g/L Tryptone, 5 g/L Yeast Extract, 0.5g/L NaCl) at 28°C and 200 rpm. Bacterial cultures were resuspended to an OD_600_ of 0.05 and mixed 1:1 with VdAve1 (± salts) in 96-well plates, yielding final conditions of OD_600_ = 0.025, 8 µM VdAve1, and 0, 6.25, 12.5, 25, 50 mM NaCl, CaCl_2_, or MgCl_2_. For lipoteichoic acid (LTA) inhibition assays, VdAve1 (240 µg/mL) was preincubated with purified *B. subtilis* LTA (240 µg/mL; Sigma-Aldrich, St. Louis, MO, USA) for 30 min at room temperature prior to mixing 1:1 with a *B. subtilis* culture with an OD_600_ of 0.05. Growth was monitored by OD_600_ measurements over time using a CLARIOstar® plate reader (BMG LABTECH, Ortenberg, Germany) at 22⍰°C with shaking before each read for 10 seconds at 300 rpm.

VdAve1-derived peptides (GenScript; Proteogenix) were tested under similar conditions. *B. subtilis* and *Sphingopyxis macrogoltabida* were grown in low-salt ¼ LB or tryptic soy broth, respectively, resuspended to OD_600_ = 0.05, and incubated with 50 µM peptides in 96-well plates. Growth was monitored as described above at 25⍰°C. Peptide net charge was calculated as the sum of ionizable groups using an online tool (https://www.bachem.com/knowledge-center/peptide-calculator/).

### VdAve1 activity assay on fungal protoplasts

Protoplasts from the fungal species *Trichoderma viride* and *Monilinia fructicola* were obtained as previously described (55). Protoplasts were pelleted at 100 x g for 1 min and resuspended in 1 M sorbitol and 10 mM MOPS at pH 6.3 to a final concentration of 10^5^-10^6^ protoplasts/mL. Protoplasts were incubated with 8 µM of VdAve1, Triton X-100 or sterile ultrapure water. Following 30 minutes of incubation, intact protoplasts were quantified for the different treatments using a hemocytometer.

### Membrane potential measurements

*B. subtilis* membrane potential measurements were performed as described previously (27). *B. subtilis* cultures were grown at 30°C in 0.5x YT, until OD_600_ of 0.1. Cultures were supplemented with 8, 4 or 2 μM VdAve1 (1, 0.5 and 0.25x MIC) or 2 μg/mL Gramicidin.

### Transcriptome analysis

B. subtilis strain 168 was grown in low-salt LB medium at 30°C while shaking at 200 rpm. Flasks containing 10 mL fresh medium were inoculated to an OD_600_ of 0.05. Cells were grown until OD_600_ of 0.3 was reached and then treated with 4 μM of VdAve1 or sterile ultrapure water. Samples were collected after 5 and 20 min of exposure. Total RNA was extracted, DNase-treated, and purified. Ribosomal RNA was depleted and sequencing libraries were prepared. The libraries were sequenced on an Illumina NextSeq platform, and differential gene expression analysis was performed using DESeq2 within the Galaxy platform. Detailed procedures are provided in SI Materials and Methods.

### Transposon sequencing (Tn-seq)

A genome-wide transposon insertion library in *B. subtilis* was generated based on established protocols (56, 57), with minor modifications. The library was propagated under control conditions or in the presence of 4 μM VdAve1 for approximately nine generations. Genomic DNA was isolated and processed to enrich for transposon-genome junctions, followed by Illumina sequencing. Sequencing reads were mapped to the *B. subtilis* reference genome, and insertion frequencies at TA dinucleotide sites were quantified to assess gene-level fitness contributions. Differences in insertion abundance between conditions were evaluated statistically using unpaired two-sided Student’s t-tests. Detailed procedures are provided in SI Materials and Methods.

### Digestion of *B. subtilis* peptidoglycan with VdAve1 and cellosyl

Peptidoglycan from *B. subtilis* bFB66 was prepared as previously published (58). VdAVe1 activity was assayed with PG or muropeptides (generated by overnight digestion with the muramidase cellosyl) as substrate. Reactions (50 μL) containing 10 μM VdAve1 were performed under two buffer conditions to account for salt sensitivity: 20 mM HEPES/NaOH (pH 7.5) with 50 mM NaCl and 1 mM MgCl_2_, or 20 mM HEPES/NaOH (pH 7.5) without added salt. Control samples contained no VdAve1. Samples were incubated at 37°C for 16 h with shaking, terminated by boiling. The PG samples were acidified to pH 4.8 and digested with muramidase cellosyl (0.5 mg/ml) at 37°C for 16 h with shaking, the reaction was then stopped by boiling for 10 min. The samples with muropeptide substrates and the cellosyl digested PG samples were reduced with sodium borohydride, and acidified (pH 3.5–4.0). Muropeptides were analysed by HPLC as described previously (39). Two control samples were taken. One bFB66 PG sample was digested with cellosyl and the other with VdAVe1, in either of the two buffers and processed as described above.

## Data availability

The sequencing data have been deposited in the European Nucleotide Archive (ENA) under accession number PRJEB38296.

## ACKNOWLEDGEMENTS

B.P.H.J.T. acknowledges funding by the Alexander von Humboldt Foundation in the framework of an Alexander von Humboldt Professorship endowed by the German Federal Ministry of Education and Research and is furthermore supported by DFG under Germany’s Excellence Strategy – EXC 2048/1 – Project ID: 390686111 and by the DFG – Project ID 458090666 / CRC1535/1. W.V. was supported by the UK BBSRC (BB/W013630/1).

## AUTHOR CONTRIBUTIONS

G.C.P., N.C.S. and B.P.H.J.T. conceived the project. G.C.P., N.C.S., A.K.R, W.V., L.H., J.R.M., D.F., A.M. and B.P.H.J.T designed the experiments. G.C.P., N.C.S., A.K.R, B.W., K.K., T.S., J.B., D.F. and A.M. carried out the experiments. G.C.P., N.C.S., A.K.R, W.V., L.H., J.R.M., D.F., A.M. and B.P.H.J.T analysed the data. G.C.P., N.C.S. and B.P.H.J.T. wrote the manuscript. All authors read and approved the final manuscript.

