## Supplemental Information for "A fungal pathogen effector that shapes host plant microbiota kills bacteria through lipoteichoic acid binding and membrane disruption"

### SUPPORTING INFORMATION

#### SI Materials and Methods

##### VdAve1 protein production and purification

Recombinant VdAve1 was produced as inclusion bodies in *E. coli* BL21 cells and purified under denaturing conditions by $Ni^{2+}$ affinity chromatography using an ÄKTA go system (Cytiva, Marlborough, MA, USA). Purified protein was refolded by stepwise dialysis using a redox-assisted glutathione system with decreasing concentrations of guanidinium chloride, as previously described (1). Final buffer exchange was performed into 30 mM potassium phosphate buffer (pH 6.5).

For VdAve1 isotopic labelling, *E. coli* BL21 cells were grown in M9 minimal medium prepared according to a published EMBL-EBI protocol (Heidelberg, Germany). For $^{15}N$ labelling, the medium was supplemented with 0.5 g/L $^{15}N$ $H_4Cl$ (Sigma-Aldrich, St. Louis, USA). For $^{13}C$, $^{15}N$ labelling, 4 g/L D-Glucose-$^{13}C_6$ (Deutero, Kastellaun, Germany) was used as the sole carbon source in addition to $^{15}N$ $H_4Cl$. Production, purification and refolding were performed as described above.

##### Sample preparation and NMR spectroscopy

NMR samples for structure determination, including all 3D experiments (see Table S 1), and $^1H,^{15}N$ TRACT experiments, were prepared in 1.7 mm NMR tubes with a final volume of 40 µL. Spectra were acquired with a Bruker Avance III HD 600 MHz spectrometer equipped with TCI Microcryoprobe$^{TM}$. Titrations were performed in 5 mm NMR tubes at 600 µL final sample volume either in a Bruker Avance III HD 600 MHz or a Bruker Avance III HDX 500 MHz spectrometer equipped with a TCI cryogenic probe. All experiments were acquired at 298 K, unless otherwise stated. NMR spectra were processed with NMRPipe (2) or TopSpin 3.6 (Bruker, Billerica, USA) and analysed using CcpNmr 3.1 software suit (3).

Samples for 3D experiments contained 600-650 µM [$^{13}C$, $^{15}N$] VdAve1. For $^1H$, $^{15}N$-TRACT experiments, samples were prepared at 200, 400, 600 and 766 µM VdAve1. LTA and peptidoglycan were titrated over samples containing 150 µM VdAve1 concentration. To reduce potential oxidized cysteines, $^1H$, $^{15}N$ HSQC spectra of VdAve1 were acquired with 4 mM β-mercaptoethanol. All samples were prepared in 30 mM phosphate buffer (pH 6.6) containing 8% $D_2O$, 200 µM DSS-$d_6$, and 0.01% $NaN_3$.

**Spectral assignment and structure calculation**

Backbone and side chain resonances were assigned using 2D $^1$H-$^{15}$N HSQC, $^1$H-$^{13}$C HSQC, 3D HNCO, HN(CA)CO, HNCA, CBCA(CO)NH, HNCACB, HN(CO)CA, HCC(CO)NH, HBHA(CO)NH, CC(CO)NH, HNHA, HCCH-TOCSY, $^1$H-$^{15}$N TOCSY-HSQC, $^1$H-$^{15}$N NOESY-HSQC with 90, 100 and 150 ms mixing times, $^1$H-$^{13}$C NOESY-HSQC with 200 ms mixing time. For more details see Table S 1. All figures showing structures were generated with PyMOL (The PyMOL Molecular Graphics System, Version 3.0 Schrödinger, LLC). The APBS Electrostatics Plugin 3.4.1 was used to visualize surface charge distribution. The structural homology search was performed with Foldseek (4).

**$^1$H,$^{15}$N TRACT experiments**

$^1$H,$^{15}$N TRACT experiments were performed to estimate rotational correlation times ($\tau_c$). A series of 30 relaxation delays was recorded (in s): 0, 0.0015, 0.003, 0.0045, 0.006, 0.0075, 0.009, 0.0105, 0.012, 0.0135, 0.015, 0.01875, 0.0225, 0.02625, 0.03, 0.0375, 0.045, 0.0525, 0.060, 0.0675, 0.075, 0.09, 0.1125, 0.135, 0.15, 0.3, 0.6, 0.75, 1.125 and 1.5. Experimental parameters are summarized in Table S1.

**RNA extraction and processing**

For *B. subtilis* RNA isolation, 0.5 mL of culture was rapidly transferred into 2 mL screw-cap tubes containing 1.5 g of 0.1 mm glass beads, 500 µL phenol–chloroform–isoamyl alcohol (25:24:1), and 50 µL of 10% SDS. Samples were immediately mixed and snap-frozen in liquid nitrogen. Cells were disrupted by bead beating in a TissueLyser (10 cycles of 30 s at 6,000 rpm). Following lysis, samples were centrifuged at 13,000 × g for 5 min and the aqueous phase was transferred to a fresh tube containing 400 µL chloroform. After mixing and centrifugation (13,000 × g for 5 min), the aqueous phase was recovered and RNA was precipitated using ethanol and sodium acetate. RNA pellets were collected by centrifugation (30 min at 13,000 × g), resuspended in nuclease-free water, and treated with DNase prior to downstream processing.

**rRNA depletion, library preparation, and sequencing**

Ribosomal RNA was removed using the MICROBExpress™ kit according to the manufacturer's instructions. RNA-seq libraries were generated using the NEBNext® Ultra™ II Directional RNA Library Prep Kit for Illumina® following the manufacturer's protocol. Libraries were sequenced on an Illumina NextSeq 550 instrument using a NextSeq 500/550 High Output v2.5 kit to generate 75-bp reads. Raw sequencing reads were processed and analysed using the Galaxy platform. Differential gene expression analysis was conducted using DESeq2 (Galaxy version 2.11.40.6+galaxy1) with default parameters unless otherwise specified.

**Transposon insertion reaction and library construction**

In vitro transposition reactions contained 6 µg genomic DNA from *B. subtilis* strain BSB1, 2 µg pCJ4 transposon plasmid, and 0.4 µg Himar1-C9 transposase in a buffer consisting of 9.5% glycerol, 93.5 mM NaCl, 9.5 mM $MgCl_2$, 3.6 µM BSA, 1.9 mM dithiothreitol, and 20.5 mM HEPES (pH 7.9). Reactions were incubated overnight at 30°C. DNA was precipitated, resuspended in 10 mM $MgCl_2$, 1 mM dithiothreitol, and 50 mM Tris-HCl (pH 7.8). To repair transposon insertion junctions, 4 µL of 2.5 mM dNTPs and 0.5 µL of T4 DNA Polymerase (New England Biolabs, Ipswich, MA, USA) were added, and the reaction was incubated at 12°C for 20 minutes. Following heat inactivation, 0.2 µL of 2.6 mM NAD and 0.5 µL of *E. coli* DNA ligase were added, and the mixture was incubated overnight at 16°C to repair nicked DNA strands. Transposon-tagged genomic DNA was transformed into chemically competent *B. subtilis* BSB1 cells, and transformants (approximately $2.5 \times 10^5$) were selected on agar plates before pooling and storing at −80°C.

**Tn-seq experiment**

The transposon library pre-culture was grown in low-salt LB medium (10 g/L tryptone, 5 g/L yeast extract, 0.5 g/L NaCl) at 28°C. Cultures were started at $OD_{600}$ = 0.1 and propagated in triplicate flasks under control or VdAve1-supplemented conditions (4 µM). Serial passaging was performed by transferring 0.75 mL at mid-log phase ($OD_{600} \approx 0.8$), into fresh medium to $OD_{600}$ = 0.1 maintaining selective pressure across three passages.

Genomic DNA was digested with MmeI (New England Biolabs, Ipswich, MA, USA) and treated with calf intestinal phosphatase prior to purification. A custom Illumina R1 adapter was annealed from oligonucleotides BW-TnR1-S and BW-TnR1-AntiS (Table S10) and ligated using T4 DNA ligase. Adapter-ligated fragments were purified using AMPure XP beads (Beckman Coulter, Indianapolis, IN, USA). PCR amplification was performed with a universal primer (BW-TnPCR-Uni) and barcoded primers (Table S10). Libraries were size-selected (~165 bp) on 8% polyacrylamide gels and eluted in 0.1× TE buffer.

**Sequencing and bioinformatic processing**

Libraries were sequenced on an Illumina NextSeq 550 platform using a High Output v2.5 kit (75-cycle single-end run). Data processing was conducted using Python and the web-based Galaxy platform. Adapter trimming was performed with Trimmomatic (Galaxy v0.36.5), and transposon inverted repeat sequences were removed using Cutadapt (Galaxy v1.16.5). Reads of 13–18 bp were retained for mapping. Reads were aligned to the *B. subtilis* genome (NC_000913) using Bowtie2 (Galaxy v2.3.4.3). A customized *map_functions* identified 183,963 TA dinucleotide insertion sites across coding sequences and quantified read abundance per site. Across libraries, approximately 9.53 million reads were obtained per condition, covering 112,457 TA sites in 4,280 genes, with an average of 85 reads per insertion site. Expected insertion counts per gene were estimated based on the number of TA sites per coding sequence. Observed counts were normalized to expected values to generate gene-level insertion ratios. Differential abundance between conditions was assessed using unpaired two-sided Student's t-tests.

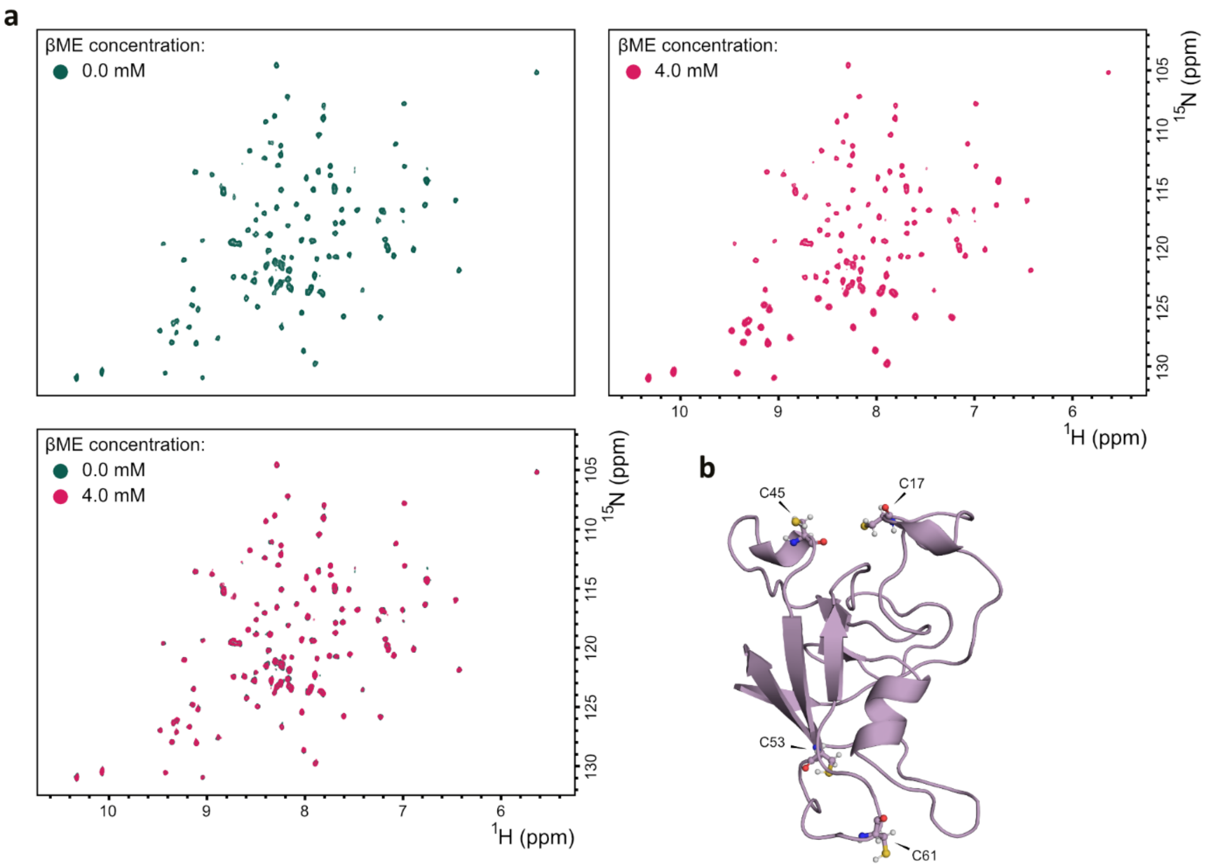


**Figure S1. VdAve1 cysteines do not form disulphide bridges. a.** $^{1}$H,$^{15}$N HSQC spectra of VdAve1 (green) and VdAve1 with the reducing agent β-mercaptoethanol (βME) (pink). Individual spectra and their superposition are shown. **b.** NMR-solved VdAve1 structure showing the side chains of the cysteines as stick and balls.

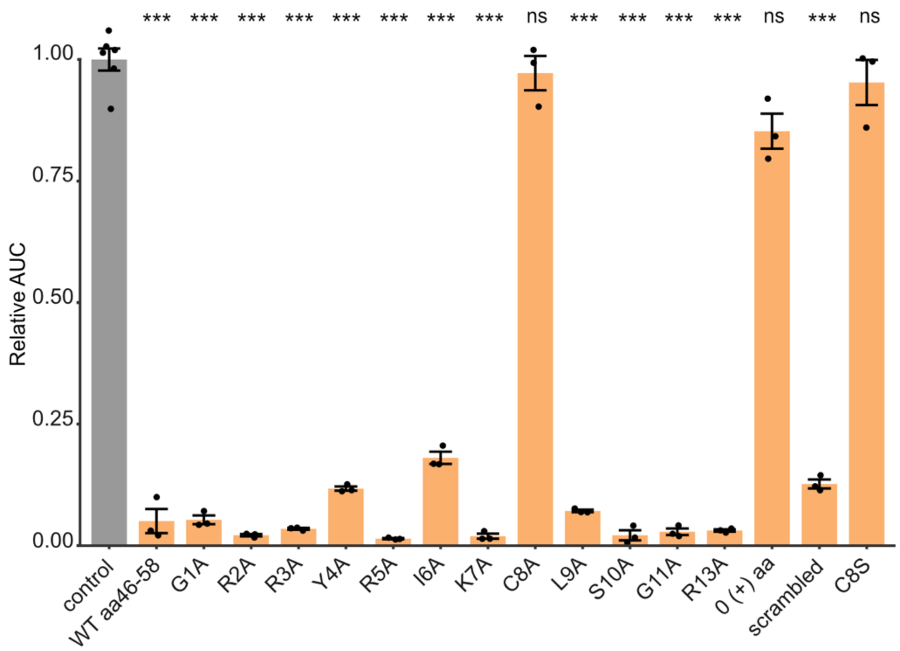


**Figure S2. VdAve1 derived peptide activity against *Sphingopyxis macrogoltabida.*** Antimicrobial activity test of peptide aa46-58 and its mutants. The plot shows area under the growth curve (AUC) of *S. macrogoltabida* treated with the respective peptide relative to the control treatment. Bars indicate mean, error bars represent standard error, individual data points are shown (n=6 for the control, n=3 for all others). Statistical significance was assessed by one-way ANOVA and post-hoc Tukey HSD test (****P* ≤0.001; ** *P* ≤0.001; * *P* ≤0.05; ns, not significant).

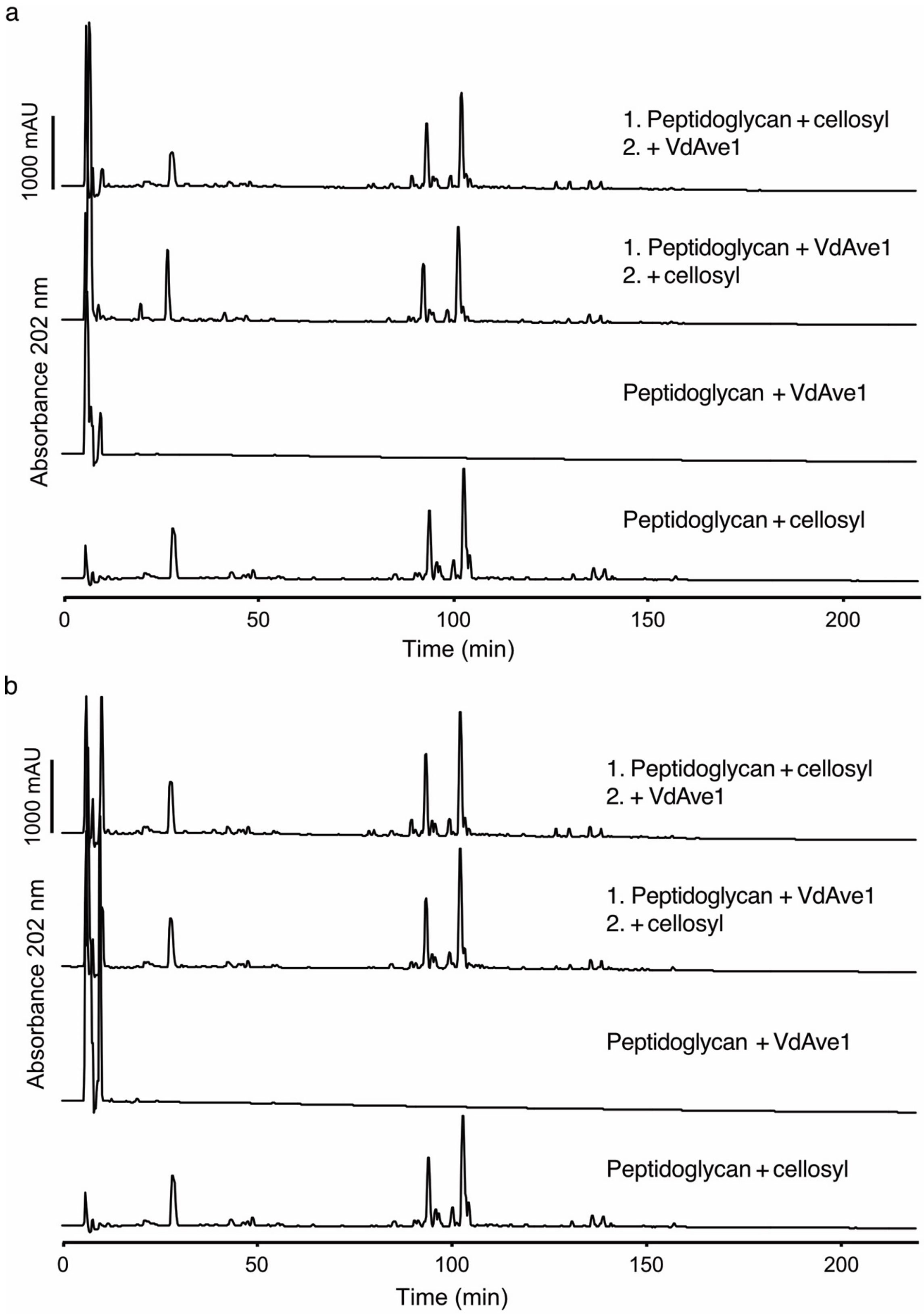


**Figure S3. VdAve1 does not hydrolyse peptidoglycan or muropeptides.** HPLC analyses of peptidoglycan (PG) from *Bacillus subtilis* incubated overnight with VdAve1. The muramidase cellosyl released disaccharide peptide subunits (muropeptides), which were analysed by HPLC. **a.** Analysis of VdAve1 activity in 20 mM HEPES/NaOH (pH 7.5) with 50 mM NaCl and 1 mM $MgCl_2$ buffer with *B. subtilis* bFB66 muropeptide or peptidoglycan as substrate. **b.** Analysis of VdAve1 activity in 20 mM HEPES/NaOH (pH 7.5) (with no additional salt) and *B. subtilis* bFB66 muropeptide or peptidoglycan as substrate.

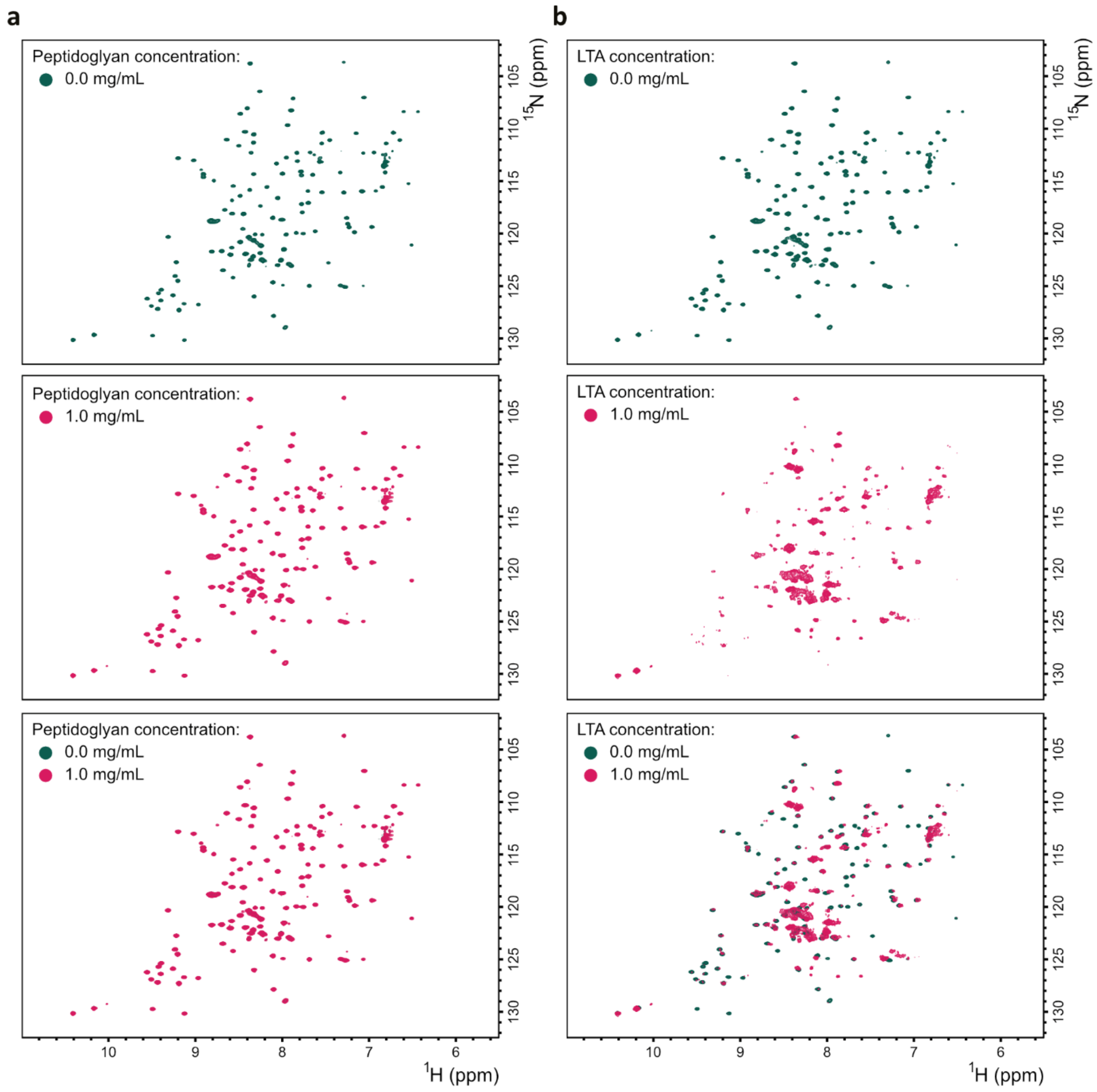


**Figure S4. VdAve1 interactions with *Bacillus subtilis* cell wall components. a.** $^{1}$H,$^{15}$N HSQC spectra of VdAve1 (green) and VdAve1 with 1 mg/mL peptidoglycan (pink). Individual spectra and their superposition are shown. **b.** $^{1}$H,$^{15}$N HSQC spectra of VdAve1 (green) and VdAve1 with 1 mg/mL peptidoglycan (pink). Individual spectra and their superposition are shown.

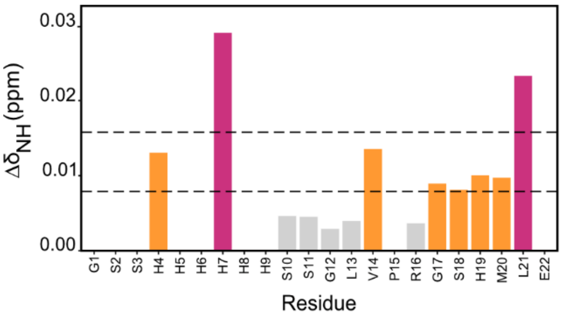


**Figure S5. Chemical shift perturbances of the protein His-tag.** The $\Delta\delta_{NH}$ values were color-coded, $\Delta\delta_{NH}>2\sigma$ pink and $\Delta\delta_{NH}>\sigma$ orange.

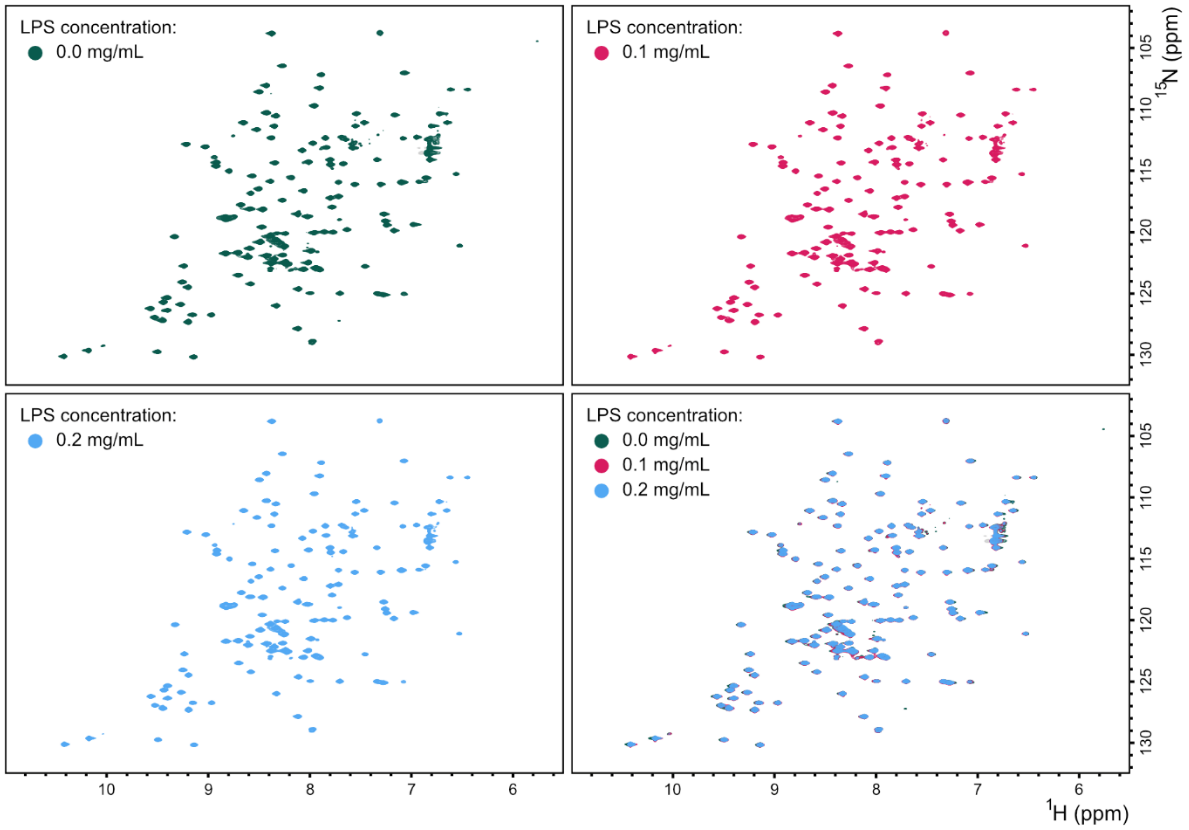


**Figure S6. VdAve1 does not bind lipopolysaccharide (LPS) from *Pseudomonas aeruginosa*. a.** $^{1}$H,$^{15}$N HSQC spectra of VdAve1 with different concentrations of LPS: 0.0 mg/mL (green), 0.1 mg/mL (pink), 0.2 mg/mL (cyan). Individual spectra and their superposition are shown.

#### Table S1. Details on experimental set-up of NMR experiments

| Exp. | Name from Bruker library | NS | SW* (ppm) | TD* | O1* (ppm) | D1 (s) | Comments |
|---|---|---|---|---|---|---|---|
| $^{1}H,^{15}N$ HSQC | trosyetf3gpsi.2 | 4-16 | 32/16 | 128/2048 | 117/4.68 | 1.5 | J=90 Hz |
| $^{1}H,^{13}C$ HSQC | hsqcetgpsp.2 | 8 | 70/16 | 128/2048 | 40/4.68 | 1 | J=145 Hz |
| HNCO | hncogpwg3d | 16 | 14/29/16 | 128/80/2048 | 173.5/116.5/4.68 | 1 | NUS 40% |
| HN(CA)CO | hncacogpwg3d | 16 | 14/29/16 | 128/80/2048 | 173.5/116.5/4.68 | 1 | NUS 40% |
| HNCA | hncagpwg3d | 16 | 30/29/16 | 128/80/2048 | 53.2/116.5/4.68 | 1 | NUS 40% |
| HNCACB | hncacbgpwg3d | 104 | 69/29/14 | 128/64/2048 | 42/116.5/4.68 | 1 | NUS 40% |
| CBCA(CO)NH | cbcaconhgpwg3d | 32 | 69/29/14 | 128/64/2048 | 42/116.5/4.68 | 1 | NUS 40% |
| HN(CO)CA | hncocagpwg3d | 16 | 30/29/16 | 128/80/2048 | 53.2/116.5/4.68 | 1 | NUS 40% |
| HCC(CO)NH | hccconhgpwg3d3 | 64 | 16/29/16 | 128/64/2048 | 4.68/116.5/4.68 | 1.5 | NUS 40% |
| HBHA(CO)NH | hbhaconhgpwg3d | 56 | 16/29/14 | 128/64/2048 | 4.68/116.5/4.68 | 1 | NUS 40% |
| CC(CO)NH | ccconhgp3d | 48 | 80.8/29/14 | 128/64/2048 | 43/116.5/4.68 | 1 | NUS 40% |
| HNHA | hnhagp3d | 16 | 30/14/14 | 64/128/2048 | 115/4.68/4.68 | 1 | NUS 405 |
| HCCH-TOCSY | hcchdigp3d2 | 32 | 80.8/80.8/14 | 96/96/2048 | 40/40/4.7 | 1 | - |
| $^{1}H$-$^{15}N$ TOCSY-HSQC | mlevhsqcetf3gp3d | 32 | 16/29/16 | 128/80/2048 | 4.68/116.5/4.68 | 2 | NUS 43% |
| $^{1}H$-$^{15}N$ NOESY-HSQC | noesyhsqcetf3gp3d | 16 | 16/29/14 | 128/80/2048 | 4.68/116.5/4.68 | 1.5 | NUS 53%, 100 ms and 150 ms |
| $^{1}H$-$^{15}N$ NOESY-HSQC | noesyhsqcetf3gp3d | 16 | 16/29/14 | 128/80/2048 | 4.68/116.5/4.68 | 1.5 | 90 ms |
| $^{1}H$-$^{13}C$ NOESY-HSQC | noesyhsqcetgp3d | 32 | 14/80/14 | 128/80/2048 | 4.7/43/4.7 | 1.2 | 200 ms |
| $^{1}H,^{15}N$-TRACT | 15n1h-tract-alpha, 15n1h-tract-beta | 40 | 33.3/16 | 30/2048 | 98.65/4.68 | 2 | - |

* F1/F2/F3

**Table S2. Structure calculation statistics**

| **NMR distances and constraints** | |
|---|---|
| Total number of distance restraints | 1643 |
| Number of intraresidual restraints (\|*i*-*j*\| = 0) | 355 |
| Number of sequential restraints (\|*i*-*j*\| = 1) | 516 |
| Number of medium-range restraints (1 < \|*i*-*j*\| < 5) | 202 |
| Number of long-range restraints (\|*i*-*j*\| ≥ 5) | 570 |
| Number of torsion angle restraints | 182 |
| **Structure statistics** | |
| Distance restraint violations > 0.1 Å | 8 ± 3 |
| Max. distance restraint violation (Å) | 0.26 ± 0.04 |
| Angle restraint violations > 5.0° | 7 ± 1 |
| Maximal angle restraint violation (°) | 19.10 ± 0.52 |
| Residues in most favored Ramachandran plot regions | 81.50% |

**Table S3. Backbone torsion angle predictions obtained using TALOS-N**

| Residue | ϕ (°) | ψ (°) | Δϕ (°) | Δψ (°) | Prediction distance | S² | Database matches | Chemical shifts | Classification |
|---|---|---|---|---|---|---|---|---|---|
| 1_D | 9.999.000 | 9.999.000 | 0.000 | 0.000 | 0.000 | 0.000 | 0 | 12 | None |
| 2_L | -120.323 | 145.502 | 11.784 | 12.366 | 0.564 | 0.848 | 25 | 17 | Strong |
| 3_G | -146.917 | 165.337 | 19.033 | 15.956 | 0.369 | 0.881 | 25 | 17 | Strong |
| 4_T | -137.492 | 153.519 | 10.462 | 6.052 | 0.362 | 0.909 | 25 | 17 | Strong |
| 5_A | -135.105 | 151.880 | 10.173 | 9.411 | 0.375 | 0.926 | 25 | 18 | Strong |
| 6_S | -128.877 | 150.107 | 17.341 | 12.289 | 0.490 | 0.930 | 25 | 18 | Strong |
| 7_Y | -136.466 | 153.690 | 13.322 | 10.233 | 0.628 | 0.924 | 25 | 18 | Strong |
| 8_Y | -134.332 | 147.412 | 16.961 | 14.229 | 0.807 | 0.901 | 25 | 18 | Strong |
| 9_N | -121.650 | 89.210 | 18.818 | 19.845 | 0.959 | 0.859 | 25 | 15 | Bad |
| 10_P | -67.655 | 153.638 | 8.742 | 11.682 | -0.107 | 0.844 | 25 | 13 | Strong |
| 11_P | -69.908 | 156.328 | 9.515 | 10.813 | 1.033 | 0.845 | 25 | 13 | Strong |
| 12_Y | -110.368 | 154.380 | 29.214 | 10.035 | 1.109 | 0.868 | 7 | 16 | Warn |
| 13_L | -117.589 | 135.617 | 21.499 | 21.243 | 1.052 | 0.885 | 25 | 16 | Strong |
| 14_P | -70.246 | 146.417 | 10.918 | 10.439 | 0.722 | 0.887 | 25 | 16 | Strong |
| 15_T | -124.311 | 168.485 | 11.481 | 7.154 | 0.586 | 0.900 | 25 | 16 | Strong |
| 16_A | -66.326 | -22.448 | 6.047 | 7.193 | 0.737 | 0.876 | 25 | 18 | Strong |
| 17_C | -82.503 | -8.223 | 10.807 | 7.169 | 0.758 | 0.867 | 25 | 17 | Strong |
| 18_G | 92.062 | 3.726 | 11.386 | 14.153 | 1.012 | 0.819 | 7 | 16 | Warn |
| 19_G | 84.649 | 11.679 | 15.072 | 18.517 | 0.778 | 0.805 | 7 | 16 | Warn |
| 20_S | -87.445 | -8.658 | 9.632 | 12.703 | 0.525 | 0.762 | 10 | 17 | Generous |
| 21_N | -81.046 | 131.737 | 26.102 | 15.440 | 0.411 | 0.736 | 25 | 16 | Strong |
| 22_P | -61.132 | -26.351 | 5.276 | 7.131 | 0.341 | 0.720 | 25 | 16 | Strong |
| 23_S | -70.697 | -16.546 | 5.511 | 5.338 | 0.306 | 0.722 | 25 | 16 | Strong |
| 24_Q | -90.829 | -6.790 | 8.588 | 6.147 | 0.297 | 0.747 | 25 | 18 | Strong |
| 25_F | -79.747 | 143.497 | 11.181 | 13.146 | 0.232 | 0.730 | 25 | 16 | Strong |
| 26_P | -68.990 | 153.859 | 7.995 | 9.645 | 0.252 | 0.727 | 25 | 16 | Strong |
| 27_S | -64.025 | 137.836 | 7.164 | 6.248 | 0.382 | 0.718 | 25 | 15 | Strong |
| 28_G | 92.213 | -7.995 | 8.389 | 9.740 | 0.600 | 0.754 | 9 | 17 | Warn |
| 29_N | -80.895 | 130.734 | 15.588 | 10.697 | 0.839 | 0.813 | 3 | 17 | Warn |
| 30_L | -85.940 | 127.200 | 10.946 | 30.319 | 0.800 | 0.884 | 25 | 18 | Strong |
| 31_F | -142.497 | 159.101 | 12.234 | 7.915 | 0.714 | 0.933 | 25 | 18 | Strong |
| 32_V | -130.291 | 150.560 | 10.395 | 12.278 | 0.845 | 0.947 | 25 | 18 | Strong |
| 33_A | -103.479 | 130.565 | 17.718 | 13.572 | 1.169 | 0.927 | 25 | 18 | Strong |
| 34_V | -116.449 | 156.525 | 22.204 | 11.772 | -2.597 | 0.904 | 25 | 18 | Strong |
| 35_S | 62.973 | 37.472 | 7.776 | 11.190 | -2.495 | 0.861 | 10 | 18 | Bad |
| 36_D | -100.316 | -2.708 | 16.570 | 14.983 | -4.696 | 0.848 | 10 | 17 | Generous |
| 37_G | 85.354 | 10.906 | 12.047 | 16.427 | 12.886 | 0.821 | 25 | 17 | Strong |
| 38_L | -103.302 | 141.951 | 26.304 | 12.764 | 2.284 | 0.827 | 10 | 17 | Generous |

| | | | | | | | | | |
|---|---|---|---|---|---|---|---|---|---|
| 39_W | -86.827 | 144.671 | 28.748 | 14.744 | 1.592 | 0.820 | 10 | 18 | Generous |
| 40_D | -61.816 | -39.086 | 12.219 | 7.256 | 0.000 | 0.837 | 10 | 18 | Bad |
| 41_N | -73.390 | -23.218 | 16.344 | 15.258 | 0.000 | 0.814 | 10 | 16 | Bad |
| 42_G | 91.539 | 0.989 | 9.148 | 10.804 | 0.922 | 0.819 | 25 | 16 | Strong |
| 43_A | -65.694 | -30.586 | 5.846 | 10.083 | 0.885 | 0.826 | 25 | 16 | Strong |
| 44_A | -76.980 | -19.574 | 11.882 | 9.777 | 1.116 | 0.858 | 10 | 18 | Generous |
| 45_C | -69.830 | -27.512 | 8.926 | 10.700 | 0.000 | 0.844 | 10 | 17 | Bad |
| 46_G | 111.342 | -24.839 | 50.365 | 45.421 | 0.673 | 0.826 | 25 | 17 | Warn |
| 47_R | -69.395 | 144.217 | 5.994 | 8.905 | 0.476 | 0.839 | 25 | 17 | Strong |
| 48_R | -120.502 | 144.005 | 13.210 | 8.384 | 0.275 | 0.877 | 25 | 18 | Strong |
| 49_Y | -136.375 | 151.318 | 8.517 | 9.085 | 0.185 | 0.916 | 25 | 18 | Strong |
| 50_R | -111.835 | 131.353 | 12.140 | 6.868 | 0.230 | 0.919 | 25 | 18 | Strong |
| 51_I | -125.159 | 142.744 | 7.926 | 8.768 | 0.260 | 0.912 | 25 | 18 | Strong |
| 52_K | -132.323 | 142.831 | 9.224 | 11.667 | 0.349 | 0.883 | 25 | 18 | Strong |
| 53_C | -130.927 | 135.080 | 18.090 | 10.943 | 0.557 | 0.848 | 25 | 18 | Strong |
| 54_L | -101.850 | 131.725 | 16.097 | 9.511 | 0.727 | 0.785 | 4 | 18 | Warn |
| 55_S | -162.501 | 170.260 | 9.525 | 10.708 | 0.767 | 0.750 | 25 | 16 | Bad |
| 56_G | 107.193 | 178.410 | 40.145 | 26.804 | 1.013 | 0.658 | 4 | 16 | Warn |
| 57_A | -63.882 | -32.597 | 5.854 | 8.705 | 0.850 | 0.636 | 25 | 16 | Strong |
| 58_R | -73.574 | -27.453 | 19.476 | 41.841 | 1.262 | 0.615 | 3 | 16 | Warn |
| 59_G | -67.287 | -33.573 | 9.246 | 10.049 | 1.373 | 0.712 | 7 | 16 | Warn |
| 60_S | -65.895 | -29.087 | 5.346 | 7.903 | 1.443 | 0.789 | 25 | 16 | Strong |
| 61_C | -93.429 | 0.686 | 14.637 | 17.829 | 2.251 | 0.874 | 25 | 18 | Strong |
| 62_K | -94.687 | -2.633 | 14.906 | 12.233 | 1.619 | 0.843 | 25 | 18 | Bad |
| 63_D | -102.407 | 0.158 | 14.159 | 12.786 | 1.215 | 0.834 | 5 | 17 | Warn |
| 64_G | 83.677 | 7.872 | 11.229 | 10.732 | 1.008 | 0.842 | 25 | 17 | Bad |
| 65_M | -119.997 | 154.938 | 14.021 | 10.156 | 0.518 | 0.887 | 25 | 17 | Strong |
| 66_I | -136.397 | 150.170 | 8.797 | 10.280 | 0.440 | 0.916 | 25 | 18 | Strong |
| 67_D | -111.012 | 137.942 | 23.160 | 15.324 | 0.527 | 0.924 | 25 | 18 | Strong |
| 68_V | -133.006 | 158.890 | 6.281 | 8.438 | 0.394 | 0.929 | 25 | 18 | Strong |
| 69_R | -119.289 | 135.911 | 12.616 | 5.589 | 0.438 | 0.925 | 25 | 18 | Strong |
| 70_V | -69.705 | 129.404 | 5.964 | 6.932 | 0.536 | 0.919 | 25 | 18 | Strong |
| 71_V | -124.185 | 150.051 | 13.963 | 18.039 | 0.758 | 0.915 | 6 | 18 | Warn |
| 72_D | -147.572 | 152.749 | 9.544 | 13.514 | 0.833 | 0.906 | 25 | 18 | Strong |
| 73_R | -82.838 | 130.508 | 21.180 | 18.513 | 1.033 | 0.899 | 25 | 18 | Strong |
| 74_A | -60.392 | -28.730 | 6.164 | 6.017 | 1.048 | 0.890 | 25 | 18 | Strong |
| 75_K | -81.025 | -7.714 | 10.648 | 9.518 | 1.228 | 0.878 | 25 | 18 | Strong |
| 76_T | -82.725 | 123.670 | 24.012 | 30.643 | 1.399 | 0.866 | 10 | 18 | Generous |
| 77_T | -80.513 | 153.203 | 27.271 | 21.742 | 1.292 | 0.833 | 8 | 18 | Warn |
| 78_V | -64.067 | -27.407 | 5.625 | 6.807 | 1.010 | 0.801 | 25 | 18 | Strong |
| 79_T | -91.597 | 2.290 | 10.800 | 12.642 | 0.820 | 0.720 | 25 | 18 | Strong |
| 80_K | -102.412 | -0.554 | 15.327 | 15.236 | 0.786 | 0.691 | 4 | 18 | Warn |

| | | | | | | | | | |
|---|---|---|---|---|---|---|---|---|---|
| 81_A | 120.081 | 46.112 | 71.762 | 55.737 | 0.718 | 0.694 | 5 | 18 | Warn |
| 82_A | -66.742 | -28.732 | 6.794 | 5.301 | 0.670 | 0.748 | 25 | 18 | Strong |
| 83_H | -141.330 | 156.625 | 13.244 | 10.709 | 0.732 | 0.785 | 9 | 18 | Warn |
| 84_K | -69.323 | 139.097 | 11.608 | 10.144 | 0.826 | 0.811 | 7 | 18 | Warn |
| 85_A | -100.170 | 149.565 | 28.709 | 8.864 | 0.759 | 0.851 | 25 | 18 | Strong |
| 86_T | -72.750 | -36.067 | 13.338 | 10.328 | 1.141 | 0.891 | 25 | 18 | Strong |
| 87_M | -143.734 | 149.294 | 12.548 | 13.362 | 0.958 | 0.898 | 25 | 18 | Strong |
| 88_I | -90.143 | 113.670 | 16.319 | 19.554 | 0.878 | 0.871 | 25 | 18 | Strong |
| 89_L | -81.914 | 143.567 | 10.967 | 13.665 | 0.797 | 0.848 | 8 | 18 | Warn |
| 90_S | -106.669 | 162.425 | 43.789 | 9.740 | 0.399 | 0.861 | 25 | 18 | Warn |
| 91_Q | -60.318 | -39.533 | 6.274 | 5.677 | 0.262 | 0.892 | 25 | 18 | Strong |
| 92_D | -64.209 | -41.207 | 3.247 | 4.224 | 0.199 | 0.922 | 25 | 18 | Strong |
| 93_S | -65.217 | -44.145 | 2.296 | 3.253 | 0.150 | 0.916 | 25 | 18 | Strong |
| 94_Y | -61.700 | -43.818 | 3.649 | 5.089 | 0.166 | 0.907 | 25 | 18 | Strong |
| 95_D | -66.553 | -25.987 | 4.293 | 8.941 | 0.181 | 0.893 | 25 | 18 | Strong |
| 96_A | -77.805 | -25.046 | 5.003 | 9.649 | 0.232 | 0.878 | 25 | 18 | Strong |
| 97_I | -112.219 | -14.200 | 9.583 | 11.668 | 0.354 | 0.867 | 25 | 18 | Strong |
| 98_V | -110.642 | 128.137 | 10.628 | 12.807 | 0.567 | 0.831 | 25 | 18 | Strong |
| 99_N | -104.929 | 147.041 | 35.568 | 22.274 | 1.501 | 0.797 | 5 | 18 | Warn |
| 100_Q | -122.793 | 134.377 | 26.475 | 15.276 | 1.071 | 0.770 | 9 | 18 | Warn |
| 101_W | -136.302 | 116.071 | 13.459 | 7.177 | 0.367 | 0.761 | 10 | 18 | Generous |
| 102_K | 54.047 | 41.764 | 5.853 | 6.983 | 0.250 | 0.772 | 25 | 16 | Strong |
| 103_G | 82.880 | 3.749 | 5.621 | 6.340 | 0.266 | 0.796 | 25 | 16 | Strong |
| 104_T | -123.728 | 150.762 | 17.450 | 20.024 | 0.440 | 0.848 | 25 | 16 | Strong |
| 105_R | -80.409 | 137.936 | 16.438 | 17.886 | 0.797 | 0.893 | 25 | 18 | Strong |
| 106_H | -74.792 | 146.005 | 15.569 | 12.996 | 0.795 | 0.919 | 10 | 18 | Generous |
| 107_K | -71.719 | -35.155 | 7.466 | 8.678 | 0.590 | 0.918 | 25 | 18 | Strong |
| 108_A | -151.493 | 159.323 | 12.412 | 7.001 | 0.578 | 0.911 | 25 | 18 | Strong |
| 109_V | -137.830 | 152.144 | 8.772 | 6.096 | 0.456 | 0.907 | 25 | 18 | Strong |
| 110_N | -135.851 | 159.946 | 28.060 | 22.210 | 0.409 | 0.908 | 25 | 18 | Bad |
| 111_I | -134.972 | 154.410 | 10.948 | 7.331 | 0.301 | 0.921 | 25 | 18 | Strong |
| 112_E | -132.795 | 150.419 | 11.342 | 13.620 | 0.244 | 0.929 | 25 | 18 | Strong |
| 113_F | -136.537 | 154.497 | 10.103 | 6.931 | 0.289 | 0.927 | 25 | 18 | Strong |
| 114_R | -140.256 | 145.062 | 12.174 | 13.425 | 0.338 | 0.902 | 25 | 18 | Strong |
| 115_Q | -72.980 | 132.875 | 9.514 | 10.973 | 0.563 | 0.861 | 25 | 18 | Strong |
| 116_I | 9.999.000 | 9.999.000 | 0.000 | 0.000 | 0.000 | 0.000 | 0 | 12 | None |

**Table S4. Structural homologs of VdAve1 identified by Foldseek**

| Description | Organism | Seq. Id. | E-Value | Probability |
|---|---|---|---|---|
| Crystal structure of apo-clavibacter Michiganensis expansin | *Clavibacter michiganensis* | 19.4 | 2.41E-04 | 1.00 |
| Crystal structure of d78n mutant clavibacter michiganensis expansin in complex with cellohexaose | *Clavibacter michiganensis subsp. michiganensis NCPPB 382* | 16.6 | 2.26E-04 | 1.00 |
| Crystal structure of cotton alpha-like expansin GhEXLA1 | *Gossypium hirsutum* | 20.1 | 2.29E-03 | 1.00 |
| Crystal structure of EXPB1 (Zea m 1), a beta-expansin and group-1 pollen allergen from maize | *Zea mays* | 16.9 | 1.28E-03 | 1.00 |
| Structure of the Ustilago maydis chorismate mutase 1 in complex with a Zea mays kiwellin | *Zea mays* | 16.4 | 2.14E-03 | 1.00 |
| Structure of the Ustilago maydis chorismate mutase 1 in complex with KWL1-b from Zea mays | *Zea mays* | 16.7 | 1.06E-03 | 1.00 |
| Crystal structure of the kiwifruit allergen Act d 5 | *Actinidia deliciosa* | 16.1 | 1.77E-03 | 1.00 |
| Crystal structure of cerato-platanin 3 from M. perniciosa (MpCP3) | *Moniliophthora perniciosa* | 16.6 | 1.89E-03 | 1.00 |
| Crystal structure of kiwellin | *Actinidia chinensis* | 15.6 | 2.96E-03 | 1.00 |
| Crystal structure of cerato-platanin 2 from M. perniciosa (MpCP2) | *Moniliophthora perniciosa* | 17.1 | 4.64E-03 | 1.00 |
| Crystal Structure of Phl p 1, a Major Timothy Grass Pollen Allergen | *Phleum pratense* | 15.8 | 9.41E-03 | 0.99 |
| Crystal structure of the cerato-platanin-like protein Cpl1 from Ustilago maydis | *Ustilago maydis 521* | 16.3 | 8.27E-03 | 0.97 |

**Table S5. List of peptides used in this study.**

| ID | Sequence |
|---|---|
| aa1-20 | DLGTASYYNPPYLPTACGGS |
| aa11-30 | PYLPTACGGSNPSQFPSGNL |
| aa21-40 | NPSQFPSGNLFVAVSDGLWD |
| aa31-50 | FVAVSDGLWDNGAACGRRYR |
| aa41-60 | NGAACGRRYRIKCLSGARGS |
| aa51-70 | IKCLSGARGSCKDGMIDVRV |
| aa61-80 | CKDGMIDVRVVDRAKTTVTK |
| aa71-90 | VDRAKTTVTKAAHKATMILS |
| aa81-100 | AAHKATMILSQDSYDAIVNQ |
| aa91-110 | QDSYDAIVNQWKGTRHKAVN |
| aa101-116 | WKGTRHKAVNIEFRQI |
| aa43-58 | AACGRRYRIKCLSGAR |
| aa46-58 | GRRYRIKCLSGAR |
| aa45-54 | CGRRYRIKCL |
| aa45-53 | CGRRYRIKC |
| aa47-52 | RRYRIK |
| G1A | ARRYRIKCLSGAR |
| R2A | GARYRIKCLSGAR |

| ID | Sequence |
|---|---|
| R3A | GRAYRIKCLSGAR |
| Y4A | GRRARIKCLSGAR |
| R5A | GRRYAIKCLSGAR |
| I6A | GRRYRAKCLSGAR |
| K7A | GRRYRIACLSGAR |
| C8A | GRRYRIKALSGAR |
| L9A | GRRYRIKCASGAR |
| S10A | GRRYRIKCLAGAR |
| G11A | GRRYRIKCLSAAR |
| R13A | GRRYRIKCLSGAA |
| 5 (+) aa | GRRYRIKCLSGAR |
| 4 (+) aa | GRRYRIACLSGAR |
| 3 (+) aa | GRRYAIACLSGAR |
| 2 (+) aa | GRAYAIACLSGAR |
| 1 (+) aa | GAAYAIACLSGAR |
| 0 (+) aa | GAAYAIACLSGAA |
| (+) aa swapped | GKKYKIRCLSGAK |
| scrambled | RSRCIGLGRKAYR |

**Table S6. The ten most strongly induced *B. subtilis* genes in response to 0.5x MIC of VdAve1.**

| | 5 minutes | | 20 minutes | |
|---|---|---|---|---|
| Gene | Log2FC VdAve1/MQ | Adjusted p value | Log2FC VdAve1/MQ | Adjusted p value |
| *sigV* | 5.04 | 9.08E-150 | 6.19 | 5.97E-186 |
| *rsiV* | 4.72 | 2.20E-112 | 5.67 | 5.09E-211 |
| *oatA* | 4.26 | 8.48E-213 | 5.35 | 0.00E+00 |
| *yrhK* | 4.21 | 1.72E-52 | 4.72 | 8.74E-108 |
| *ymaE* | 3.84 | 7.59E-75 | 2.65 | 1.38E-23 |
| *ymzB* | 3.7 | 3.45E-60 | 2.36 | 1.78E-13 |
| *ydbO* | 2.11 | 2.21E-22 | 2.33 | 1.40E-72 |
| *maeA* | 1.65 | 6.71E-06 | 2.01 | 3.22E-22 |
| *mmgD* | 1.51 | 1.09E-14 | 1.99 | 7.47E-09 |
| *bcrC* | 1.5 | 1.08E-10 | 1.99 | 2.40E-49 |

**Table S7. Selected differentially expressed *B. subtilis* genes upon incubation with 0.5x MIC of VdAve1.**

| | | 5 minutes | | 20 minutes | |
|---|---|---|---|---|---|
| | Gene | Log2FC VdAve1/MQ | Adjusted p value | Log2FC VdAve1/MQ | Adjusted p value |
| *SigV* operon | *sigV* | 4.72 | 2.20E-112 | 6.19 | 5.97E-186 |
| | *rsiV* | 5.04 | 9.08E-150 | 5.67 | 5.09E-211 |
| | *oatA* | 4.26 | 8.48E-213 | 5.35 | 0.00E+00 |
| | *yrhK* | 3.70 | 3.45E-60 | 4.72 | 8.74E-108 |
| Cell wall metabolism | *cwlO* | -1.96 | 9.98E-55 | 0.15 | 0.57 |
| | *lytE* | -1.26 | 2.87E-22 | 0.32 | 6.53E-02 |
| | *iseA* | 4.21 | 1.72E-52 | 1.95 | 5.58E-11 |
| Teichoic acid synthesis or modification | *dltA* | 0.34 | 2.79E-03 | 0.85 | 4.17E-20 |
| | *dltB* | 0.23 | 0.29 | 0.8 | 1.34E-09 |
| | *dltC* | 0.16 | 0.47 | 0.77 | 9.05E-11 |
| | *dltD* | 0.33 | 5.52E-03 | 0.80 | 3.02E-17 |
| | *dltE* | 0.32 | 4.19E-02 | 0.85 | 4.33E-13 |
| | *tagT* | 0.68 | 1.86E-09 | 0.75 | 1.77E-11 |
| | *ugtP* | 0.33 | 4.54E-02 | 0.32 | 3.82E-02 |
| | *yfnI* | 0.53 | 2.42E-06 | 0.47 | 2.68E-05 |
| Cation export/uptake | *cadA* | 0.81 | 3.85E-06 | 0.18 | 0.57 |
| | *ydbO* | 0.90 | 2.51E-10 | 2.33 | 1.40E-72 |
| | *ykkD* | 0.81 | 9.26E-03 | 0.35 | 0.45 |
| | *nhaK* | -0.07 | 0.91 | 1.01 | 9.53E-07 |
| Membrane fluidity | *des* | -1.15 | 2.46E-06 | 0.02 | 0.98 |

**Table S8. Genes affecting *B. subtilis* sensitivity to VdAve1 identified by transposon sequencing.**

| | Gene | Relative fitness VdAve1/MQ | Adjusted p value |
|---|---|---|---|
| *SigV* operon | *rsiV* | 2.61 | 2.53E-18 |
| | *oatA* | 0.68 | 1.75E-05 |
| Cell wall metabolism | *pbpX* | 1.90E-02 | 1.56E-235 |
| | *cwlO* | 2.66E-03 | 9.25E-94 |
| | *dacA* | 0.16 | 1.10E-26 |
| Teichoic acid synthesis or modification | *dltA* | 1.76E-03 | 1.74E-292 |
| | *dltB* | 9.86E-03 | 0.00E+00 |
| | *dltC* | 2.52E-03 | 2.47E-65 |
| | *dltD* | 2.59E-03 | 0.00E+00 |
| | *dltE* | 0.27 | 1.77E-24 |
| | *galE* | 8.72E-02 | 5.29E-150 |
| | *gtaB* | 0.24 | 1.60E-21 |
| | *ugtP* | 0.12 | 4.25E-45 |
| Other genes | *greA* | 0.16 | 3.41E-18 |
| | *miaA* | 0.31 | 4.70E-08 |
| | *rsiX* | 2.36 | 1.06E-16 |

**Table S9. $^{15}N$ and $^{1}H$ chemical shit perturbances of VdAve1 in presence of 0.6 mg/mL of *Bacillus subtilis* lipoteichoic acid (LTA).**

| Residue | 15N APO | 1H APO | 15N LTA | 1H LTA | CSP |
|---|---|---|---|---|---|
| TAG-4_H | 120.4523 | 8.387343 | 120.6381 | 8.368723 | 0.013143 |
| TAG-7_H | 122.4554 | 8.237324 | 122.1859 | 8.264064 | 0.029105 |
| TAG-10_S | 118.1251 | 8.447731 | 118.1044 | 8.439253 | 0.004716 |
| TAG-11_S | 115.9366 | 7.704725 | 115.8929 | 7.70177 | 0.004615 |
| TAG-12_G | 110.5392 | 8.329976 | 110.5202 | 8.33463 | 0.003008 |
| TAG-13_L | 121.5195 | 7.983124 | 121.4834 | 7.986825 | 0.004061 |
| TAG-14_V | 122.5274 | 8.012086 | 122.3928 | 8.007606 | 0.013654 |
| TAG-16_R | 121.9394 | 8.423916 | 121.9451 | 8.431334 | 0.003752 |
| TAG-17_G | 110.2877 | 8.419199 | 110.2102 | 8.428501 | 0.009035 |
| TAG-18_S | 115.5516 | 8.173207 | 115.4899 | 8.162376 | 0.008212 |
| TAG-19_H | 121.1535 | 8.239946 | 121.0956 | 8.22328 | 0.010144 |
| TAG-20_M | 120.0795 | 8.226194 | 120.1612 | 8.237116 | 0.009826 |
| TAG-21_L | 122.9937 | 8.381109 | 122.767 | 8.392446 | 0.023376 |
| 1_D | 120.8645 | 8.299526 | 120.8884 | 8.284171 | 0.00804 |
| 2_L | 118.6624 | 8.014717 | 118.534 | 7.990885 | 0.017519 |
| 3_G | 108.0174 | 8.401788 | 107.9609 | 8.395443 | 0.006487 |
| 4_T | 113.0153 | 9.023672 | 113.0175 | 9.036509 | 0.006422 |
| 5_A | 120.3038 | 9.316472 | 120.2195 | 9.302913 | 0.010822 |
| 6_S | 113.0814 | 7.826013 | 113.0603 | 7.814948 | 0.005921 |
| 7_Y | 118.0852 | 8.577985 | 118.0816 | 8.593559 | 0.007795 |
| 8_Y | 117.3648 | 8.2519 | 117.276 | 8.244358 | 0.009649 |
| 9_N | 120.0713 | 7.952252 | 120.2727 | 7.990849 | 0.027899 |
| 12_Y | 118.7541 | 8.739156 | 118.5777 | 8.713501 | 0.011323 |
| 13_L | 115.5534 | 6.846573 | 115.4912 | 6.845851 | 0.006227 |
| 15_T | 108.5629 | 8.4845 | 108.5197 | 8.471447 | 0.007826 |
| 16_A | 122.7156 | 9.220758 | 122.7039 | 9.220712 | 0.001171 |
| 17_C | 110.4272 | 7.149753 | 110.3909 | 7.143641 | 0.004744 |
| 18_G | 106.4415 | 8.25882 | 106.3823 | 8.260775 | 0.005997 |
| 19_G | 112.7424 | 7.951866 | 112.7338 | 7.953881 | 0.001324 |
| 20_S | 114.4382 | 7.778157 | 114.4382 | 7.778157 | 0 |
| 21_N | 119.3546 | 6.972481 | 119.34 | 6.969261 | 0.002175 |
| 23_S | 114.2772 | 7.996249 | 114.2526 | 7.992354 | 0.003132 |
| 24_Q | 115.2241 | 6.545453 | 115.1756 | 6.544095 | 0.004901 |
| 25_F | 119.8553 | 7.170107 | 119.8442 | 7.166179 | 0.002256 |
| 27_S | 116.8188 | 8.575994 | 116.7786 | 8.569063 | 0.005306 |
| 28_G | 113.8952 | 8.933022 | 113.9022 | 8.927551 | 0.002826 |
| 29_N | 111.3223 | 8.329757 | 111.2753 | 8.325585 | 0.005138 |
| 30_L | 121.0806 | 6.509531 | 121.0514 | 6.508832 | 0.002942 |
| 31_F | 119.0714 | 7.246307 | 119.0732 | 7.236117 | 0.005099 |
| 32_V | 112.2815 | 8.357677 | 112.1815 | 8.351965 | 0.010398 |
| 33_A | 127.1745 | 9.4377 | 127.1468 | 9.436887 | 0.002803 |

| | | | | | |
|---|---|---|---|---|---|
| 34_V | 109.6549 | 7.938729 | 109.5939 | 7.938774 | 0.006101 |
| 35_S | 117.9555 | 7.768877 | 118.0027 | 7.784731 | 0.009226 |
| 36_D | 118.1251 | 8.447731 | 118.1044 | 8.439253 | 0.004716 |
| 37_G | 119.0714 | 7.246307 | 119.0732 | 7.236117 | 0.005099 |
| 38_L | 118.4947 | 7.257652 | 118.4466 | 7.254958 | 0.00499 |
| 39_W | 119.7805 | 7.620018 | 119.7799 | 7.617937 | 0.001042 |
| 40_D | 118.1251 | 8.447731 | 118.1044 | 8.439253 | 0.004716 |
| 42_G | 114.3175 | 8.906756 | 114.3029 | 8.912121 | 0.003052 |
| 43_A | 124.9734 | 7.688246 | 124.9549 | 7.691248 | 0.002381 |
| 44_A | 113.5069 | 6.836661 | 113.4892 | 6.841147 | 0.00286 |
| 45_C | 113.5069 | 6.836661 | 113.4892 | 6.841147 | 0.00286 |
| 46_G | 111.6105 | 8.500213 | 111.6019 | 8.494053 | 0.003199 |
| 47_R | 122.1652 | 8.334223 | 122.1702 | 8.333526 | 0.000607 |
| 48_R | 121.6429 | 8.69596 | 121.633 | 8.69047 | 0.002919 |
| 49_Y | 117.7284 | 8.66195 | 117.7355 | 8.651603 | 0.005222 |
| 50_R | 124.04 | 9.239356 | 124.0688 | 9.223151 | 0.008598 |
| 51_I | 125.6775 | 9.429115 | 125.5327 | 9.416497 | 0.015795 |
| 52_K | 122.5032 | 8.361853 | 122.459 | 8.358298 | 0.004761 |
| 53_C | 126.1933 | 9.558955 | 126.221 | 9.559077 | 0.002771 |
| 54_L | 129.7234 | 9.49634 | 129.8572 | 9.514199 | 0.016092 |
| 55_S | 112.2896 | 7.830638 | 112.1613 | 7.812596 | 0.015687 |
| 56_G | 107.1118 | 7.875492 | 107.0714 | 7.868123 | 0.005467 |
| 57_A | 125.987 | 8.322918 | 125.8302 | 8.338632 | 0.017535 |
| 58_R | 123.054 | 8.178193 | 123.064 | 8.183094 | 0.002645 |
| 59_G | 112.1858 | 8.750826 | 111.8211 | 8.722055 | 0.039203 |
| 60_S | 114.3868 | 7.651039 | 114.1194 | 7.593763 | 0.039181 |
| 61_C | 118.4889 | 8.109593 | 118.4353 | 8.119616 | 0.00734 |
| 62_K | 121.8427 | 8.254483 | 121.8497 | 8.247925 | 0.003352 |
| 63_D | 120.308 | 8.383797 | 120.321 | 8.388286 | 0.002594 |
| 64_G | 108.2319 | 7.893534 | 108.2048 | 7.882684 | 0.006065 |
| 65_M | 116.6005 | 8.051265 | 116.5681 | 8.047356 | 0.003788 |
| 66_I | 114.3147 | 8.328113 | 114.3008 | 8.320794 | 0.003913 |
| 67_D | 122.9888 | 8.048386 | 122.9478 | 8.048956 | 0.004106 |
| 68_V | 112.8023 | 9.201639 | 112.7892 | 9.202627 | 0.001401 |
| 69_R | 120.0036 | 7.758462 | 119.9223 | 7.745113 | 0.010517 |
| 70_V | 125.8841 | 9.260005 | 125.8555 | 9.252807 | 0.004597 |
| 71_V | 118.8663 | 8.773732 | 118.6966 | 8.758663 | 0.018564 |
| 72_D | 117.0521 | 7.69964 | 117.0379 | 7.700476 | 0.001473 |
| 73_R | 123.4934 | 8.683381 | 123.5131 | 8.666819 | 0.008512 |
| 74_A | 127.8336 | 8.099024 | 127.8156 | 8.093438 | 0.003322 |
| 75_K | 114.9398 | 8.787368 | 114.7291 | 8.797848 | 0.021714 |
| 76_T | 103.657 | 7.289496 | 103.411 | 7.268914 | 0.026664 |
| 77_T | 115.9463 | 7.084486 | 115.9908 | 7.065962 | 0.010274 |
| 78_V | 121.3312 | 8.579588 | 121.3133 | 8.592283 | 0.006597 |

| | | | | | |
|---|---|---|---|---|---|
| 79_T | 115.4468 | 8.564992 | 115.4386 | 8.557062 | 0.004049 |
| 80_K | 122.5587 | 8.224018 | 122.6998 | 8.205716 | 0.016816 |
| 81_A | 124.6289 | 8.104524 | 124.7813 | 8.1249 | 0.018331 |
| 82_A | 123.0746 | 7.894492 | 122.99 | 7.902139 | 0.009287 |
| 83_H | 116.0393 | 7.544528 | 115.7006 | 7.534487 | 0.034234 |
| 84_K | 122.772 | 7.479459 | 122.7613 | 7.703196 | 0.111874 |
| 85_A | 125.0864 | 7.282408 | 124.9077 | 7.355078 | 0.040491 |
| 86_T | 117.1796 | 7.779485 | 117.1248 | 7.788707 | 0.007163 |
| 87_M | 114.4382 | 7.778157 | 114.4153 | 7.777955 | 0 |
| 88_I | 121.7028 | 8.811229 | 121.6388 | 8.80485 | 0.007146 |
| 89_L | 130.1603 | 9.127317 | 130.1564 | 9.133678 | 0.003203 |
| 90_S | 112.3162 | 7.067901 | 112.2572 | 7.06364 | 0.006275 |
| 91_Q | 119.9311 | 7.839288 | 119.8406 | 7.837827 | 0.009076 |
| 92_D | 113.3059 | 8.065172 | 113.279 | 8.07293 | 0.004718 |
| 93_S | 115.9366 | 7.704725 | 115.8929 | 7.70177 | 0.004615 |
| 94_Y | 124.1906 | 8.566472 | 124.1381 | 8.57255 | 0.006072 |
| 95_D | 115.8282 | 8.373174 | 115.7973 | 8.376162 | 0.003431 |
| 96_A | 119.3839 | 7.232252 | 119.3376 | 7.228434 | 0.005004 |
| 97_I | 107.0213 | 7.058408 | 106.9723 | 7.069189 | 0.00729 |
| 98_V | 122.877 | 7.923767 | 122.8604 | 7.917277 | 0.003646 |
| 99_N | 126.3631 | 9.397991 | 126.3099 | 9.406648 | 0.006861 |
| 100_Q | 120.8024 | 8.485173 | 120.8094 | 8.483699 | 0.00102 |
| 101_W | 121.9979 | 8.59456 | 122.0434 | 8.595461 | 0.004567 |
| 102_K | 126.7552 | 8.967003 | 126.5565 | 8.951941 | 0.021249 |
| 103_G | 103.7735 | 8.368874 | 103.7694 | 8.36235 | 0.003287 |
| 104_T | 114.0756 | 7.78022 | 114.2554 | 7.802649 | 0.021187 |
| 105_R | 120.6223 | 8.327704 | 120.5487 | 8.324944 | 0.007493 |
| 106_H | 125.3299 | 9.396101 | 125.0672 | 9.393698 | 0.0227 |
| 107_K | 124.4944 | 9.207678 | 124.4658 | 9.206029 | 0.002978 |
| 108_A | 116.0768 | 7.293411 | 116.1149 | 7.301206 | 0.00545 |
| 109_V | 111.0384 | 8.638703 | 110.9109 | 8.633157 | 0.013044 |
| 110_N | 118.7819 | 8.816786 | 118.7092 | 8.806259 | 0.008976 |
| 111_I | 114.5937 | 8.909597 | 114.6743 | 8.921473 | 0.01001 |
| 112_E | 119.5308 | 8.453869 | 119.3167 | 8.466539 | 0.022327 |
| 113_F | 118.8984 | 8.823867 | 118.7092 | 8.806259 | 0.020866 |
| 114_R | 116.4101 | 8.488763 | 116.3356 | 8.492281 | 0.00766 |
| 115_Q | 127.3124 | 9.188696 | 127.2965 | 9.189065 | 0.001604 |
| 116_I | 128.883 | 7.962245 | 128.8872 | 7.970267 | 0.004033 |

**Table S10. Primers used in the Tn-seq analysis.**

| Name | Name Sequence (5’ --> 3’) Application | Application |
|---|---|---|
| BW-TnR1-S | TCTTTCCCTACACGACGCTCTTCCGATCTNN | Tn-seq library preparation |
| BW-TnR1-AntiS | AGATCGGAAGAGCGTCGTGTAGGGAAAGA | Tn-seq library preparation |
| BW-TnPCR-Uni | AATGATACGGCGACCACCGAGATCTACACTCTTTCCCTACACGACGCTCTTCCGATCT | Tn-seq library preparation |
| BW-TnPCR-BC43 | CAAGCAGAAGACGGCATACGAGATGCTGTAGTGACTGGAGTTCAGACGTGTGCTCTTCCGATCAGACCGGGGACTTATCATCCAACCTGT | Tn-seq library preparation |
| BW-TnPCR-BC44 | CAAGCAGAAGACGGCATACGAGATATTATAGTGACTGGAGTTCAGACGTGTGCTCTTCCGATCAGACCGGGGACTTATCATCCAACCTGT | Tn-seq library preparation |
| BW-TnPCR-BC45 | CAAGCAGAAGACGGCATACGAGATGAATGAGTGACTGGAGTTCAGACGTGTGCTCTTCCGATCAGACCGGGGACTTATCATCCAACCTGT | Tn-seq library preparation |
| BW-TnPCR-BC46 | CAAGCAGAAGACGGCATACGAGATTCGGGAGTGACTGGAGTTCAGACGTGTGCTCTTCCGATCAGACCGGGGACTTATCATCCAACCTGT | Tn-seq library preparation |
| BW-TnPCR-BC47 | CAAGCAGAAGACGGCATACGAGATCTTCGAGTGACTGGAGTTCAGACGTGTGCTCTTCCGATCAGACCGGGGACTTATCATCCAACCTGT | Tn-seq library preparation |
| BW-TnPCR-BC48 | CAAGCAGAAGACGGCATACGAGATTGCCGAGTGACTGGAGTTCAGACGTGTGCTCTTCCGATCAGACCGGGGACTTATCATCCAACCTGT | Tn-seq library preparation |
